# Glial type specific regulation of CNS angiogenesis by HIFα-activated different signaling pathways

**DOI:** 10.1101/2020.01.31.929661

**Authors:** Sheng Zhang, Bokyung Kim, Xiaoqing Zhu, Xuehong Gui, Yan Wang, Zhaohui Lan, Preeti Prabhu, Kenneth Fond, Aijun Wang, Fuzheng Guo

**Affiliations:** Institute for Pediatric Regenerative Medicine, Shriners Hospitals for Children/UC Davis School of Medicine, CA 95817; Department of Neurology, School of Medicine, UC Davis, CA 95817; Qingdao University, Qingdao, China; Department of Surgery, School of Medicine, UC Davis, CA 95817

## Abstract

The mechanisms by which oligodendroglia modulate CNS angiogenesis remain elusive. Previous *in vitro* data suggest that oligodendroglia regulate CNS endothelial cell proliferation and blood vessel formation through hypoxia inducible factor alpha (HIFα)-activated Wnt (but not VEGF) signaling. Using *in vivo* genetic models, we show that HIFα in oligodendroglia is necessary and sufficient for angiogenesis independent of CNS regions. At the molecular level, HIFα stabilization in oligodendroglia does not perturb Wnt signaling but remarkably activates VEGF. At the functional level, genetically blocking oligodendroglia-derived VEGF but not Wnt significantly decreases oligodendroglial HIFα-regulated CNS angiogenesis. Interestingly, blocking astroglia-derived Wnt signaling reduces astroglial HIFα-regulated CNS angiogenesis. Together, our *in vivo* data clearly demonstrate that oligodendroglial HIFα regulates CNS angiogenesis through Wnt-independent and VEGF-dependent signaling. Our findings represent an alternative mechanistic understanding of CNS angiogenesis by postnatal glial cells and unveil a glial cell type-dependent HIFα-Wnt axis in regulating CNS vessel formation.

## Introduction

The vasculature of the central nervous system (CNS), which is developed exclusively through angiogenesis, plays a crucial role in providing neural cells with nutrients and oxygen. CNS angiogenesis, the growth of new blood vessels from pre-existing ones, starts during embryonic development and matures during postnatal development in human and rodent brains, for example, by the age of one month in rodents^1^. Dysregulated CNS angiogenesis negatively impacts postnatal brain development and functional recovery from brain injuries^2–4^. The current study aimed to dissect the molecular regulation of postnatal CNS angiogenesis using *in vivo* genetic animal models.

The developing CNS parenchyma is exposed to physiological hypoxia with local oxygen concentration ranging from 0.5% to 7%^5^. Hypoxia inducible factor α (HIFα) is a critical regulator that adapts neural cells to hypoxic conditions. The transcription factor HIFα, including HIF1α and HIF2α, is subjected to constant degradation. Von Hippel-Lindau (VHL), a negative regulator of HIFα’s transcriptional activity, plays an essential role in HIFα degradation. Under low oxygen or upon VHL disruption, HIF1α and HIF2α degradation is impaired and subsequently translocate into the nuclei where they regulate downstream target genes through forming transcriptional active complexes with the constitutive HIF1β^6,7^. Previous data suggest that HIFα function in neural precursor cells is required for embryonic brain vascular development^8^. Recent data including those from our own laboratory show that HIFα function in oligodendroglial lineage cells may play a pivotal role in regulating postnatal angiogenesis in the brain white matter^9^ and in the spinal cord^10^. However, the molecular mechanisms underlying oligodendroglial HIFα-regulated angiogenesis are still controversial and remain incompletely defined.

The current concept stated that oligodendroglial HIFα promotes CNS angiogenesis through activating signaling pathway of Wnt but not vascular endothelial growth factor (VEGF)^9^. However, this “Wnt-dependent” view was supported only by *in vitro* studies and pharmacological manipulations^9^, in which “pathological” activation of Wnt signaling, poor cell-type selectivity and/or off-target effects of small compounds cannot be excluded. In this study, we presented compelling *in vivo* evidence supporting an alternative view in our mechanistic understanding of oligodendroglial HIFα-regulated CNS angiogenesis. Our *in vivo* genetic knockout data reveal that oligodendroglial HIFα regulates endothelial cell proliferation and angiogenesis in a VEGF-dependent but Wnt-independent manner and this regulation is independent of CNS regions during postnatal development. Very interestingly, our data also demonstrate that postnatal astroglia regulate CNS angiogenesis at least in part through HIFα-activated Wnt signaling, unveiling a glial cell type-specific HIFα-Wnt connection in the CNS.

## Results

Endothelial cell (EC) proliferation is an essential step of angiogenesis and the blood vessel density is an end-point reflection of angiogenesis. Therefore, we used EC proliferation and vessel density as *in vivo* readouts of angiogenesis^9^. To quantify blood vessel density, we used the basement membrane marker Laminin to label blood vessels and employed a semi-automated approach to calculate the percentage of Laminin-occupying area among total assessed area (Supplementary Fig. 1). To determine whether oligodendroglial HIFα is required for angiogenesis throughout the postnatal CNS, *Cre-LoxP* approach was used to genetically ablate or stabilize HIFα and EC proliferation and vessel density were analyzed in the brain and the spinal cord.

### Oligodendroglia regulate CNS angiogenesis through HIFα-mediated signaling in a manner independent of CNS regions

We used *Cnp-Cre* line^11^ to generate *Cnp-Cre:Hif1α*^fl/fl^ (HIF1α conditional knockout, cKO), *Cnp-Cre:Hif2α*^fl/fl^ (HIF2α cKO), and *Cnp-Cre: Hif1α*^fl/fl^ *:Hif2α*^fl/fl^ (HIF1α/HIF2α or HIFα double cKO) mutants (Supplementary Fig. 2). Mice carrying Cnp-Cre transgene alone did not display any developmental abnormalities compared with non-Cre animals as previously reported^11^ and supported by our assessment of CNS angiogenesis and motor function (Supplementary Fig. 3). HIF1α cKO or HIF2α cKO did not influence blood vessel density, indicating a compensatory effect of oligodendroglial HIF1α and HIF2α on angiogenesis. In contrast, HIF1α/HIF2α double cKO (refer to as HIFα cKO hereafter) significantly impaired CNS angiogenesis evidenced by reduced blood vessel density (Fig. 1a-c) and diminished EC proliferation not only in the cerebral cortex but also in the spinal cord (Fig. 1d-g), suggesting that oligodendroglial HIFα is necessary for CNS angiogenesis.

**Figure 1.**
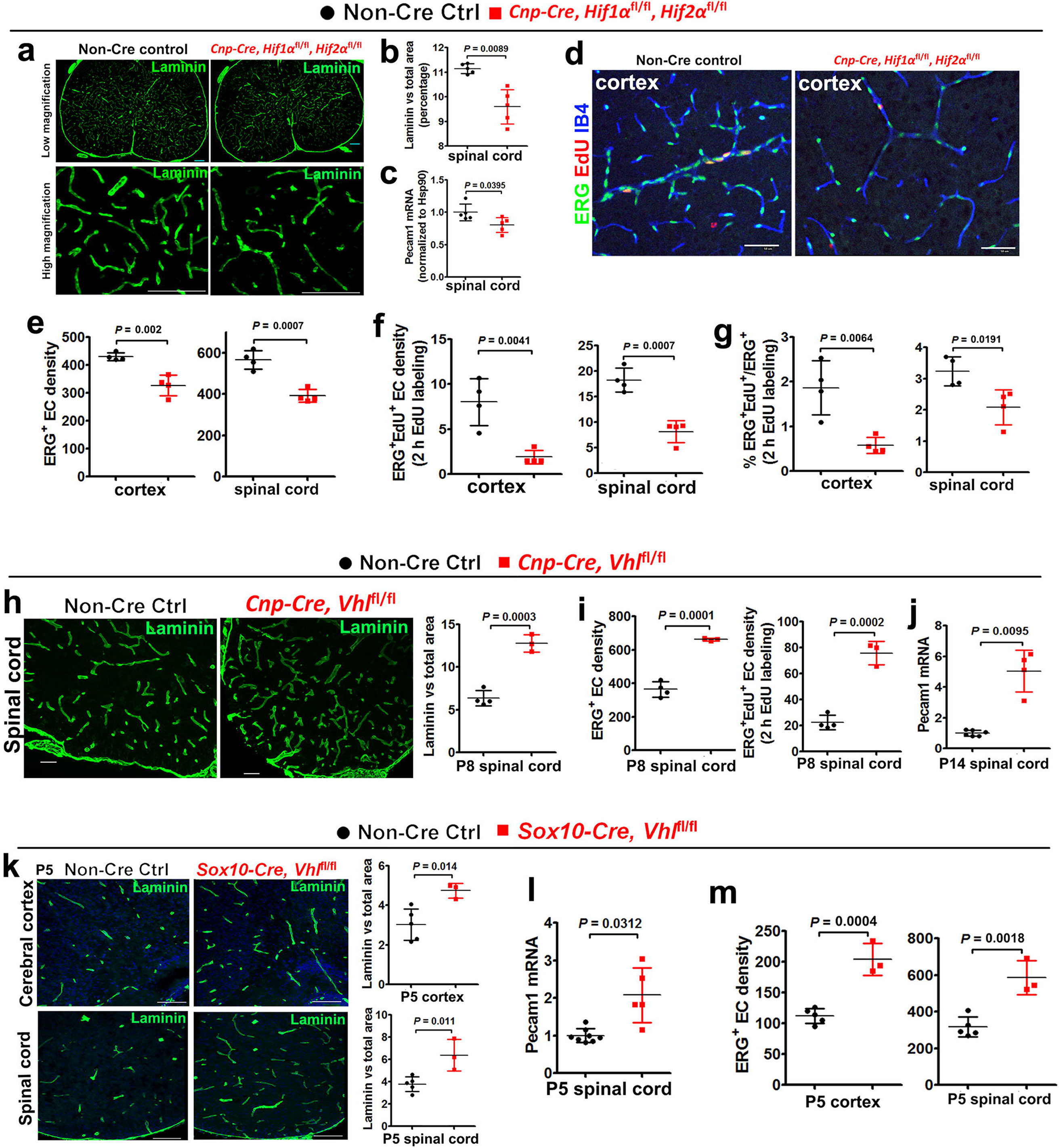
CNS region-independent regulation of angiogenesis by oligodendroglial HIFα. **a**, representative images of blood vessels labeled by the basement membrane marker Laminin in the spinal cord of *Cnp-Cre:Hif1α*^fl/fl^:*Hif2α*^fl/fl^ and non-Cre littermate controls at postnatal day 14 (P14). Scale bars = 100 μm. **b**, percentage of Laminin^+^ area among total assessed area. Two-tailed Student’s *t* test with Welch’s correction, t_(4.618)_ = 4.767. **c**, RT-qPCR assay of *Pecam1* mRNA (a.k.a. CD31), a marker of endothelial cells (ECs), in the P14 spinal cord. Two-tailed Student’s *t* test, t_(8)_ = 2.526. **d-f**, representative images of immunohistochemical staining of ERG (a nuclear marker of ECs), isolectin B4 (IB4, labeling blood vessel basement membrane), and EdU (labeling actively dividing cells) and densities of marker positive cells (# per mm^2^) in *Cnp-Cre*, *Hif1α*^fl/fl^, *Hif2α*^fl/fl^, and littermate control mice at P14. Two hours EdU pulse labeling. Two-tailed Student’s *t* test, ERG^+^, t_(6)_ = 5.200 cortex, t_(6)_ = 6.358 spinal cord; ERG^+^EdU^+^, t_(6)_ = 4.496 cortex, t_(6)_ = 6.365 spinal cord. **g**, percentage of ERG^+^ ECs that are EdU^+^. Two-tailed Student’s *t* test, t_(6)_ = 4.089 cortex, t_(6)_ = 3.181 spinal cord. **h**, representative images and quantification of Laminin in the spinal cord of P8 *Cnp-Cre, Vhl*^fl/fl^ and non-Cre control mice. Two-tailed Student’s *t* test, *t*_(5)_ = 8.831. **i**, densities (per mm^2^) of ERG^+^ ECs and ERG^+^EdU^+^ proliferating ECs in the P8 spinal cord. Two-tailed Student’s *t* test, *t*_(5)_ = 11.10 ERG^+^, *t*_(5)_ = 9.981 ERG^+^EdU^+^. **j**, RT-qPCR of *Pecam1* mRNA in P14 spinal cord. Two-tailed Student’s *t* test with Welch’s correction, *t*_(3.081)_ = 5.941. **k**, representative images and quantification of Laminin in P5 *Sox10-Cre, Vhl*^fl/fl^ and non-Cre controls. Two-tailed Student’s *t* test, *t*_(6)_ = 3.625 spinal cord, *t*_(6)_ = 3.428 cortex. **l**, RT-qPCR of *Pecam1* mRNA in P5 spinal cord. Two-tailed Student’s *t* test with Welch’s correction, *t*_(4.309)_ = 3.257. **m**, densities (per mm^2^) of ERG^+^ ECs at P5. Two-tailed Student’s *t* test, *t*_(6)_ = 6.964 cortex, *t*_(6)_ = 5.324 spinal cord. Scale bar: **a, k,** 100 μm; **d, h,** 50 μm.

The HIFα protein is constantly translated but subjected to rapid turnover via proteasome-mediated degradation, a process in which Von Hippel-Lindau product (VHL) is essential for HIFα degradation. Therefore, we employed *Cnp-Cre:Vhl*^fl/fl^ transgenic mice to genetically ablate VHL and stabilize HIFα function in oligodendroglial lineage cells (Supplementary Fig. 4). We found that the density of blood vessels and the proliferation of ECs were significantly increased in the cerebral cortex and spinal cord of *Cnp-Cre:Vhl*^fl/fl^ mice compared with those of non-Cre control mice at different time points in the early postnatal CNS (Fig. 1h-j). Stabilizing HIFα in oligodendroglial lineage cells did not have a major effect on the integrity of the blood brain (spinal cord) barrier in the adult *Cnp-Cre:Vhl*^fl/fl^ mice (Supplementary Fig. 5).

Previous studies have reported that *Cnp-Cre* primarily targets oligodendroglial lineage cells and also a subpopulation of early neural progenitor cells^12,13^. To corroborate the conclusion derived from Cnp-Cre transgenic mice, we assessed CNS angiogenesis in a different animal strain *Sox10-Cre:Vhl*^fl/fl^ in which *Sox10-Cre* mediated HIFα stabilization in the earlier stages of oligodendrocyte development in the CNS. Consistently, CNS angiogenesis was significantly increased in *Sox10-Cre:Vhl*^fl/fl^ mutants, as assessed by elevated blood vessel density (Fig. 1k), EC-specific Pecam1 mRNA expression (Fig. 1l), and EC densities (Fig. 3m). Taken together, our loss (gain)-of-function results suggest that that oligodendroglial HIFα is necessary and sufficient for angiogenesis and that the angiogenic regulation by oligodendroglial HIFα is independent of CNS regions.

### Oligodendroglial HIFα does not regulates the activity of Wnt/β-catenin signaling *in vivo* and *in vitro*

To determine whether HIFα in oligodendroglial lineage cells regulates Wnt/β-catenin signaling, we quantified the Wnt/β-catenin target gene *Axin2, Naked1*, and *Notum,* which are reliable readouts for the signaling activation^10^. We found no significant changes in the mRNA levels of those genes in HIFα-stabilized spinal cord and forebrain of *Cnp-Cre:Vhl*^fl/fl^ mutants at different time points (Fig. 2a) compared with those of non-Cre controls. Consistently, Western blot assay showed that the active form of β-catenin (dephosphorylated on Ser37 or Thr41) and Axin2 did not change (Fig. 2b), indicating that stabilizing oligodendroglial HIFα does not perturb the activity of Wnt/β-catenin signaling in the CNS. Furthermore, we found no significant change in the mRNA level of Wnt7a in the CNS of *Cnp-Cre:Vhl*^fl/fl^ mice compared with non-Cre controls (Fig. 2c). We crossed Wnt reporter transgenic mice BAT-lacZ^14^ with *Cnp-Cre:Vhl*^fl/fl^ mutants and found no difference in lacZ mRNA level in the spinal cord of BAT-lacZ/*Cnp-Cre:Vhl*^fl/fl^ mice compared with age-matched BAT-lacZ mice (Fig. 2d).

**Figure 2.**
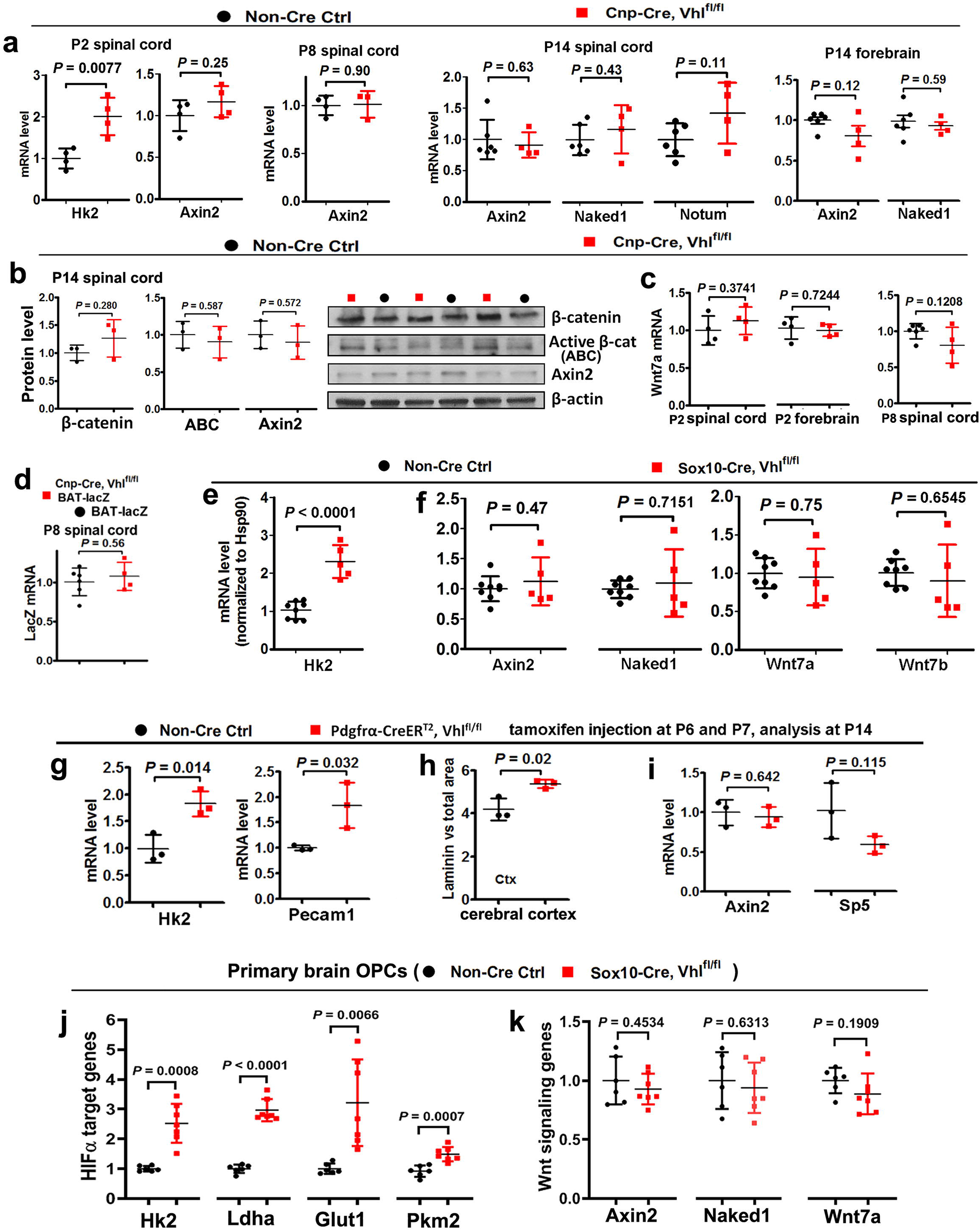
Oligodendroglial HIFα does not activate Wnt/β-catenin signaling. **a**, RT-qPCR assay of mRNA levels of HIFα target gene *Hk2* and Wnt/β-catenin target genes *Axin2, Naked1*, and *Notum*. Two-tailed Student’s *t* test, *t*_(6)_ = 3.936 *Hk2*, *t*_(6)_ = 1.275 *Axin2* at P2; *t*_(5)_ = 0.1307 *Axin2* at P8; *t*_(8)_ = 0.4906 *Axin2*, *t*_(8)_ = 0.8386 *Naked1*, *t*_(8)_ = 1.821 *Notum* at P14 spinal cord*; t*_(8)_ = 1.736 *Axin2*, *t*_(8)_ = 0.5617 *Naked1* at P14 forebrain. **b**, Western blot assay of total β-catenin, active β-catenin, and Axin2. Two-tailed Student’s *t* test, *t*_(4)_ = 1.247 β-catenin, *t*_(4)_ = 0.5869 active β-catenin, *t*_(4)_ = 0.614 Axin2. **c**, RT-qPCR of Wnt7a mRNA. Two-tailed Student’s *t* test, *t*_(6)_ = 0.960 P2 spinal cord, *t*_(6)_ = 0.3724 P2 forebrain, *t*_(8)_ = 1.736 P8 spinal cord. **d,** RT-qPCR of lacZ mRNA in Wnt/β-catenin reporter mice (BAT-lacZ) that had been crossed onto *Cnp-Cre, Vhl*^fl/fl^ and non-Cre control backgrounds. Two-tailed Student’s *t* test, *t*_(8)_ = 0.6072. **e-f**, RT-qPCR assay of mRNA level of Hk2, Axin2, Naked1, Wnt7a, and Wnt7b in the spinal cord of *Sox10-Cre:Vhl*^fl/fl^ and non-Cre controls at P5. Two-tailed Student’s *t* test, *t*_(11)_ = 7.011 *Hk2*, *t*_(11)_ = 0.7422 *Axin2*, Welch’s corrected *t*_(4.345)_ = 0.392 *Naked1*, *t*_(11)_ = 0.3216 *Wnt7a,* Welch’s corrected *t*_(4.709)_ = 0.4827 *Wnt7b.* **g,** RT-qPCR of HIFα target gene *Hk2* and EC marker *Pecam1* in P14 spinal cord. Two-tailed Student’s *t* test, *t*_(4)_ = 4.158 *Hk2*, *t*_(4)_ = 3.231 *Pecam1*. **h**, percent of laminin-occupying area among total area in the cerebral cortex at P14. Two-tailed Student’s *t* test, *t*_(4)_ = 3.724. **i**, expression of Wnt/β-catenin target gene *Axin2* and *Sp5* in the spinal cord at P14 quantified by RT-qPCR. Two-tailed Student’s *t* test, *t*_(4)_ = 0.5021 *Axin2*, *t*_(4)_ = 2.006 *Sp5.* **j-k**, RT-qPCR assay of mRNA levels of HIFα target genes and Wnt signaling genes in primary OPCs isolated from the neonatal brain of *Sox10-Cre, Vhl*^fl/fl^ and non-Cre control mice. Two-tailed Student’s *t* test, Welch’s corrected *t*_(6.282)_ = 6.065 *Hk2*, Welch’s corrected *t*_(7.898)_ = 12.92 *Ldha*, Welch’s corrected *t*_(6.199)_ = 4.009 *Glut1, t*_(11)_ = 4.654 *Pkm2*, *t*_(11)_ = 0.7772 *Axin2*, *t*_(11)_ = 0.4936 *Naked1*, *t*_(11)_ = 1.394 *Wnt7a*.

We further assessed the activity of Wnt/β-catenin signaling in a different animal strains of *Sox10-Cre:Vhl*^fl/fl^ mice. Consistent with *Cnp-Cre:Vhl*^fl/fl^ mice, HIFα was stabilized in the CNS of *Sox10-Cre:Vhl*^fl/fl^ mutants, as shown by the elevated expression of HIFα target gene *Hk2* (Fig. 2e). However, Wnt/β-catenin signaling was not perturbed, as evidenced by similar activity of Wnt/β-catenin signaling assessed at the mRNA (Fig. 2f) and protein (Supplementary Fig. 6) levels. The unperturbed activity of Wnt/β-catenin signaling was further corroborated by evidences from a time-conditional *Pdgfrα-CreER^T2^:Vhl*^fl/fl^ strain in which *Pdgfrα-CreER^T2^* elicited a greater than 85% of recombination efficiency and specificity in early postnatal oligodendrocyte progenitor cells (OPCs) (Supplementary Fig. 7). Tamoxifen-induced VHL ablation in OPCs resulted in HIFα stabilization and elevated angiogenesis, as demonstrated by significant increase in the expression of HIFα target gene *Hk2* and EC-specific *Pecam1* (Fig. 2g) and the density of cerebral blood vessels (Fig. 2h). However, the mRNA expression of Wnt target genes *Ainx2* and *Sp5* (Fig. 2i) and the protein levels of active β-catenin and Naked1 (Supplementary Fig. 8) were indistinguishable between *Pdgfrα-CreER^T2^:Vhl*^fl/fl^ mutants and non-Cre controls, indicating that Wnt/β-catenin signaling activity was not altered by oligodendroglial HIFα stabilization.

Previous study reported an autocrine activation of Wnt/β-catenin signaling in OPCs by HIFα stabilization^9^. To assess the autocrine activity of Wnt/β-catenin signaling, we treated purified primary OPCs with HIFα stabilizer DMOG^9^ in the present or absence of HIFα signaling blocker Chetomin^15^ (Supplementary Fig. 9a). Our results showed that pharmacological stabilizing HIFα activated HIFα signaling target genes (Supplementary Fig. 9b) but did not activate Wnt/β-catenin target genes nor Wnt7a and Wnt7b (Supplementary Fig. 9c) in primary OPCs isolated from neonatal brain. We also quantified the activity of Wnt/β-catenin signaling in primary OPCs which were isolated from neonatal *Sox10-Cre:Vhl*^fl/fl^ brain. Consistent with the *in vivo* data (Fig. 2e), HIFα target genes were significantly increased in primary VHL-deficient OPCs (Fig. 2j). However, neither Wnt/β-catenin target genes Axin2 and Naked1 nor Wnt7a were increased in primary VHL-deficient OPCs (Fig. 2k), suggesting that stabilizing oligodendroglial HIFα does not perturb Wnt/β-catenin signaling in primary OPCs.

To determine whether HIFα deletion affects Wnt/β-catenin signaling, we analyzed Wnt/β-catenin activity in the early postnatal CNS of Cnp-Cre:HIFα cKO and Pdgfrα-CreER^T2^:HIFα cKO mutants. Consistent with HIFα-stabilized mutants (Fig. 2), we found no evidence of Wnt/β-catenin signaling perturbation in both strains of HIFα cKO mutants (Supplementary Fig. 10). Collectively, our *in vivo* and *in vitro* data demonstrate that Wnt/β-catenin signaling is unlikely a downstream target of oligodendroglial HIFα as previously reported^9^ and suggest that oligodendroglial HIFα may regulate CNS angiogenesis independent of Wnt/β-catenin signaling.

### Blocking oligodendroglia-derived Wnt signaling does not affect HIFα-regulated CNS angiogenesis

WLS is an essential factor of Wnt secretion from Wnt-producing cells and its deficiency blocks Wnt ligands from activating the downstream pathways in Wnt-receiving cells^16–19^. To determine whether WLS deficiency affects Wnt secretion from oligodendroglial lineage cells, we knocked down WLS in primary Wnt7a-expressing OPCs and assessed Wnt secretion and autocrine Wnt/β-catenin activity (Fig. 3a, b). We expressed Wnt7a given that Wnt7a is reported as one of the Wnt ligand genes expressed in OPCs at the mRNA level^9,20^. Our enzyme-linked immunosorbent assay (ELISA) of the culture medium showed that WLS knockdown significantly reduced Wnt7a concentration secreted from Wnt7a-expressing OPCs (Fig. 3c). Autocrine Wnt/β-catenin signaling was activated in Wnt7a-expressing OPCs, as evidenced by the increased expression of Wnt target genes *Axin2* and *Sp5* (Fig. 3d), but this activation was blocked in WLS-deficient OPCs (Fig. 3d). Our data suggest that WLS is required for Wnt secretion from OPCs.

**Figure 3.**
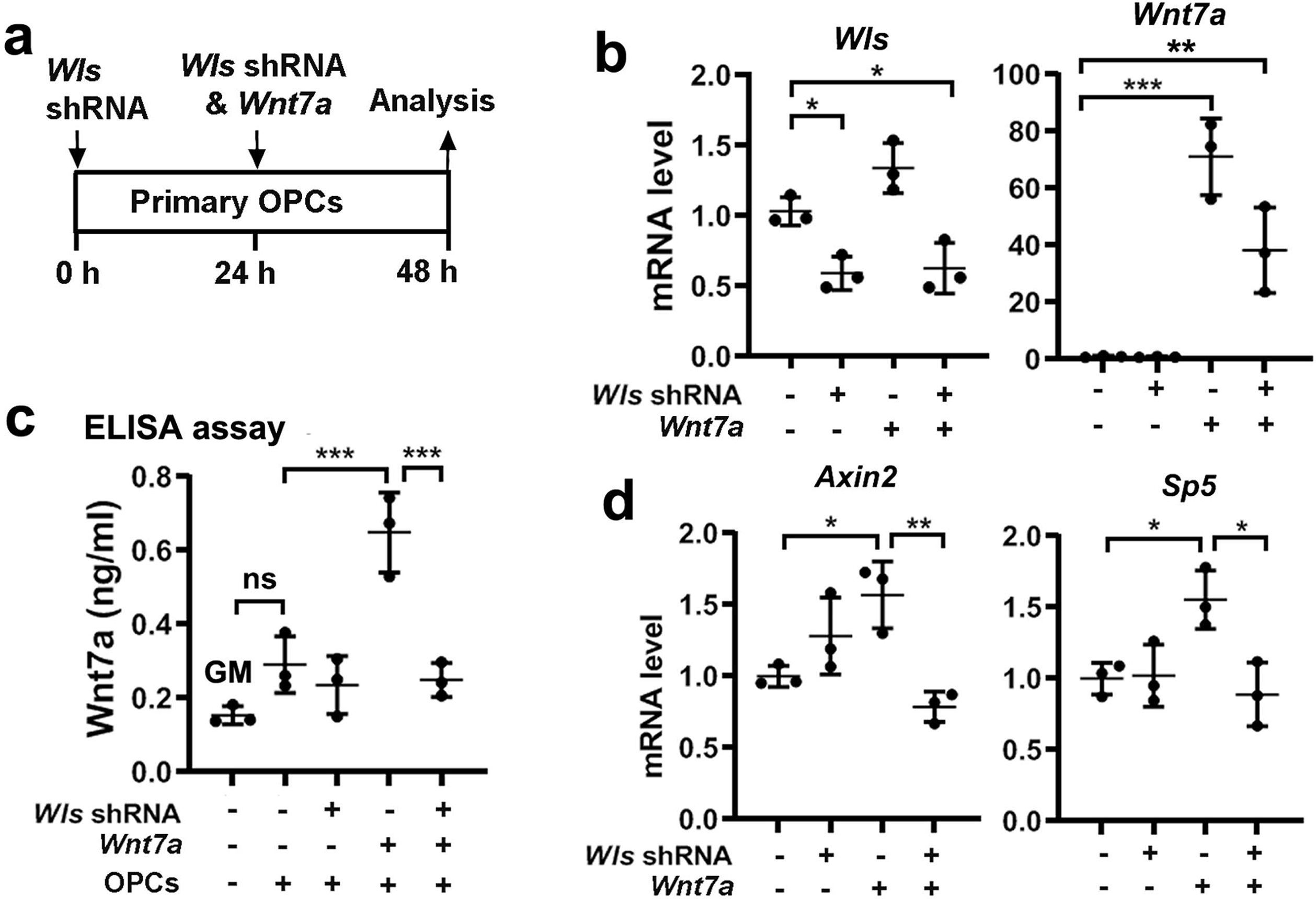
WLS is required for Wnt secretion from OPCs. **a**, brain primary OPCs growing in the growth medium (GM) were transfected with *Wls*-shRNA and *Wnt7a* plasmids, and OPC mRNA and culture medium were collected for analysis 48 hours h) after the first *Wls*-shRNA transfection. **b**, RT-qPCR assay of *Wls* and *Wnt7a* mRNA in transfected primary OPCs. One-way ANOVA followed by Tukey’s multiple comparisons, * *P* < 0.05, ** *P* < 0.01, *** *P* < 0.001. *F*_(3,8)_ = 17.41, *P* = 0.007 *Wls*, *F*_(3,8)_ = 33.82, *P* < 0.0001 *Wnt7a*. **c**, ELISA measurement of Wnt7a protein concentration in the GM in the absence and presence of OPCs in the dish with Wls-shRNA and Wnt7a transfection. One-way ANOVA followed by Tukey’s multiple comparisons, *** *P* < 0.001, ns not significant. *F*_(4, 10)_ = 21.02, *P* < 0.0001. Note that Wnt7a concentration in the GM in the presence of primary OPCs is not statistically different from that in the GM in the absence of primary OPCs. **d**, RT-qPCR assay of Wnt target genes *Axin2* and *Sp5* in OPCs. One-way ANOVA followed by Tukey’s multiple comparisons, * *P* < 0.05,** *P* < 0.01. *F*_(3, 8)_ = 9.632, *P* = 0.0049 *Axin2*, *F*_(3, 8)_ = 6.965, *P* = 0.0128 *Sp5*.

To define the putative *in vivo* role of Wnt signaling in HIFα-regulated CNS angiogenesis, we generated VHL/WLS double mutant hybrids to block Wnt secretion from HIFα-stabilized oligodendroglial lineage cells (Fig. 4a). Because constitutive *Sox10-Cre:Vhl*^fl/fl^ pups died at very early postnatal ages, we used an inducible Cre line *Sox10-CreER*^T2^ to stabilize HIFα and disrupt WLS (Fig. 4b-c) in Sox10^+^ oligodendroglial lineage cells (OPCs and differentiated oligodendroglia). Our fate-mapping data showed that *Sox10-CreER^T2^* elicited ~60% of recombination efficiency and greater than 90% of oligodendroglial specificity in Sox10^+^ oligodendroglial lineage cells in the early postnatal CNS (Supplementary Fig. 11). We confirmed that HIFα’s function was indeed stabilized in the spinal cord of *Sox10-CreER*^T2^:*Vhl*^fl/fl^ (HIFα-stabilized mice) and *Sox10-CreER*^T2^:*Vhl*^fl/fl^:*Wls*^fl/fl^(HIFα-stabilized/WLS-disrupted mice), as evidenced by the elevated expression of HIFα target gene *Hk2* (Fig. 4d) and *Ldha* (Fig. 4e) in comparison with non-Cre controls. Our analysis demonstrated that blocking Wnt secretion by disrupting WLS did not alter HIFα stabilization-elicited CNS angiogenesis in HIFα-stabilized/WLS-disrupted mice compared with HIFα-stabilized mice, which was supported by unchanged levels of endothelial *Pecam1* mRNA expression (Fig. 4f) and unchanged densities of blood vessels (Fig. 4g-j), ERG^+^ ECs (Fig. 4k), and ERG^+^EdU^+^ dividing ECs (Fig. 4l) in the spinal cord and cerebral cortex of HIFα-stabilized/WLS-disrupted mice compared with those of HIFα-stabilized mice. These data suggest that oligodendroglial lineage-derived Wnt signaling plays a minor role in HIFα-regulated angiogenesis in the early postnatal CNS.

**Figure 4.**
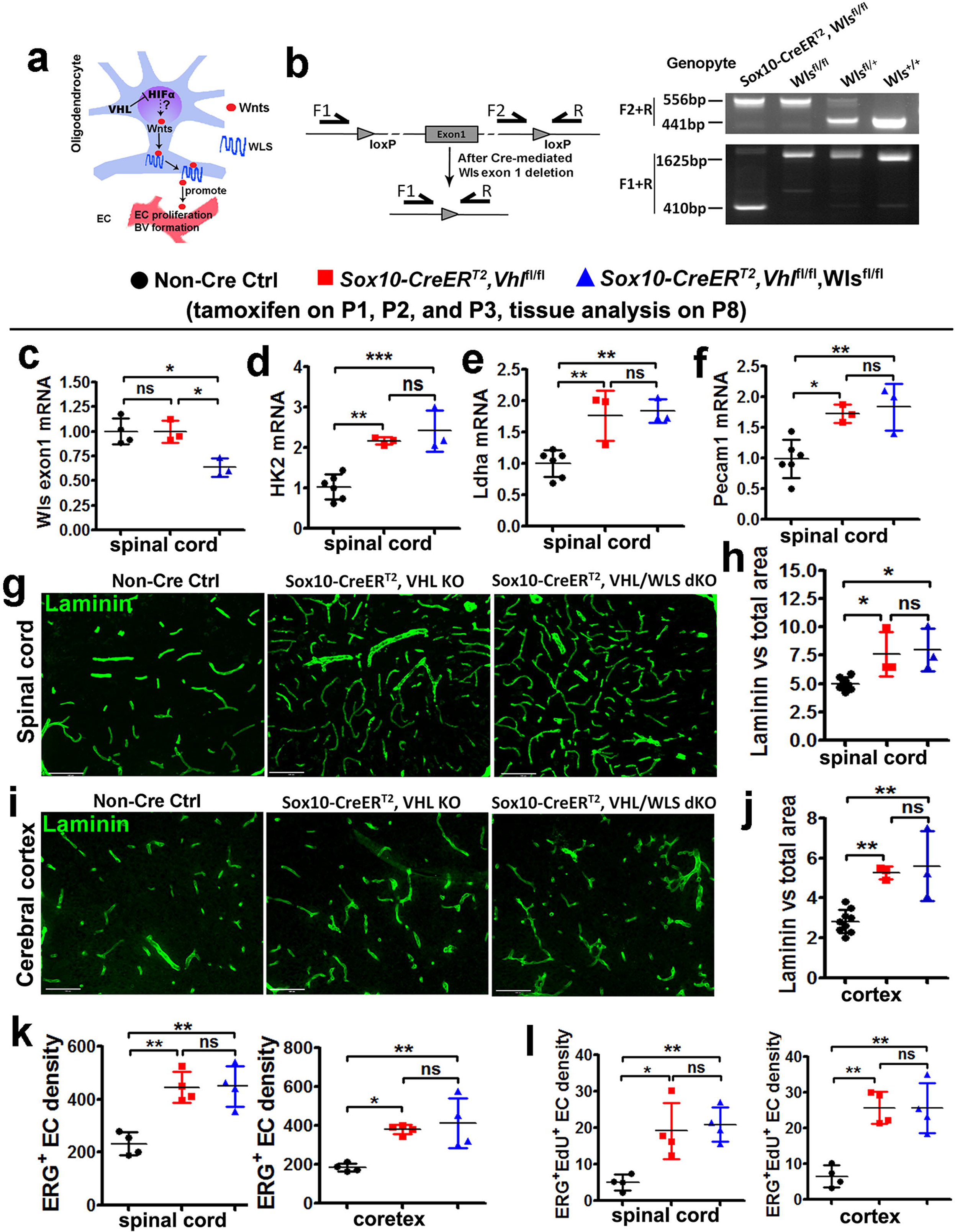
Blocking Wnt secretion from oligodendroglial lineage cells does not affect HIFα-regulated CNS angiogenesis. **a**, schematic diagram depicting putative regulation between HIFα-Wnt axis in glio-vascular units. **b,** primer design (left) and PCR detection (right) of *Wls* gene deletion. Primer pair of F2/R is for detecting *Wls* floxed allele (556bp). After Cre-mediated deletion, primer pair F1/R generates a 410bp product from the genome of *Sox10-CreER^T2^,Wls*^fl/fl^ mice. **c**, RT-qPCR assay of exon 1-coding *Wls* mRNA. One-way ANOVA followed by Tukey’s multiple comparisons, * *P* < 0.05, ns, not significant. *F*_(2, 7)_ = 10.39, *P* = 0.008. **d-f**, RT-qPCR assay of *Hk2*, *Ldha*, and *Pecam1* mRNA. One-way ANOVA followed by Tukey’s multiple comparisons, * *P* < 0.05, ** *P* < 0.01, *** *P* < 0.001, ns, not significant. *F*_(2, 9)_ = 21.73, *P* = 0.0004 *Hk2*, *F*_(2, 9)_ = 14.14, *P* = 0.0017 *Ldha*, *F*_(2, 9)_ = 10.39, *P* = 0.0046 *Pecam1*. **g-h**, representative confocal images of Laminin-positive blood vessels in the spinal cord and the percent of Laminin-positive BV area among total assessed area. One-way ANOVA followed by Tukey’s multiple comparisons, * *P* < 0.05, ns, not significant. *F*_(2, 11)_ = 8.8, *P* = 0.0052. Scale bars = 100 µm. **i-j**, representative confocal images of Laminin-positive blood vessels in the forebrain cerebral cortex and the percent of Laminin-positive BV area among total assessed area. One-way ANOVA followed by Tukey’s multiple comparisons, ** *P* < 0.01, ns, not significant. *F*_(2, 12)_ = 16.47, *P* = 0.0004. Scale bars = 100 µm. **k**, densities (#/mm^2^) of ERG^+^ endothelial cells. One-way ANOVA followed by Tukey’s multiple comparisons,* P < 0.05, ** *P* < 0.01, ns, not significant. *F*_(2, 9)_ = 16.89, *P* = 0.0009 spinal cord; Welch’s ANOVA followed by unpaired *t* test with Welch’s correction, *W*_(2, 5.346)_ = 65.98, *P* = 0.0002 cortex. **l**, densities (#/mm^2^) of ERG^+^/EdU^+^ proliferating endothelial cells (2 hrs EdU pulse labeling prior to tissue harvesting at P8). One-way ANOVA followed by Tukey’s multiple comparisons, * *P* < 0.05, ** *P* < 0.01, ns, not significant. *F*_(2, 9)_ = 10.63, *P* = 0.0043 spinal cord, *F*_(2, 9)_ = 18.43, *P* = 0.0007 cortex.

Next, we used a different Cre transgenic line *Pdgfrα-CreER*^T2^ to stabilize HIFα specifically in OPCs. OPC-specific HIFα stabilization enhanced blood vessel density (Fig. 5a-d) and increased the number of ERG^+^ ECs (Fig. 5e-f) in the spinal cord and cerebral cortex of *Pdgfrα-CreER*^T2^:*Vhl*^fl/fl^ animals compared with those in non-Cre controls. However, blocking Wnt secretion from OPCs by disrupting WLS did not affect blood vessel formation and endothelial cell density in the CNS of HIFα stabilized/WLS-disrupted mice compared with HIFα stabilized mice (*Pdgfrα-CreER*^T2^:*Vhl*^fl/fl^:*Wls*^fl/fl^ versus *Pdgfrα-CreER*^T2^:*Vhl*^fl/fl^) (Fig. 5c-f). The data from two independent Cre transgenic lines collectively suggest that oligodendroglia-derived Wnt signaling is dispensable for HIFα-regulated CNS angiogenesis *in vivo.*

**Figure 5.**
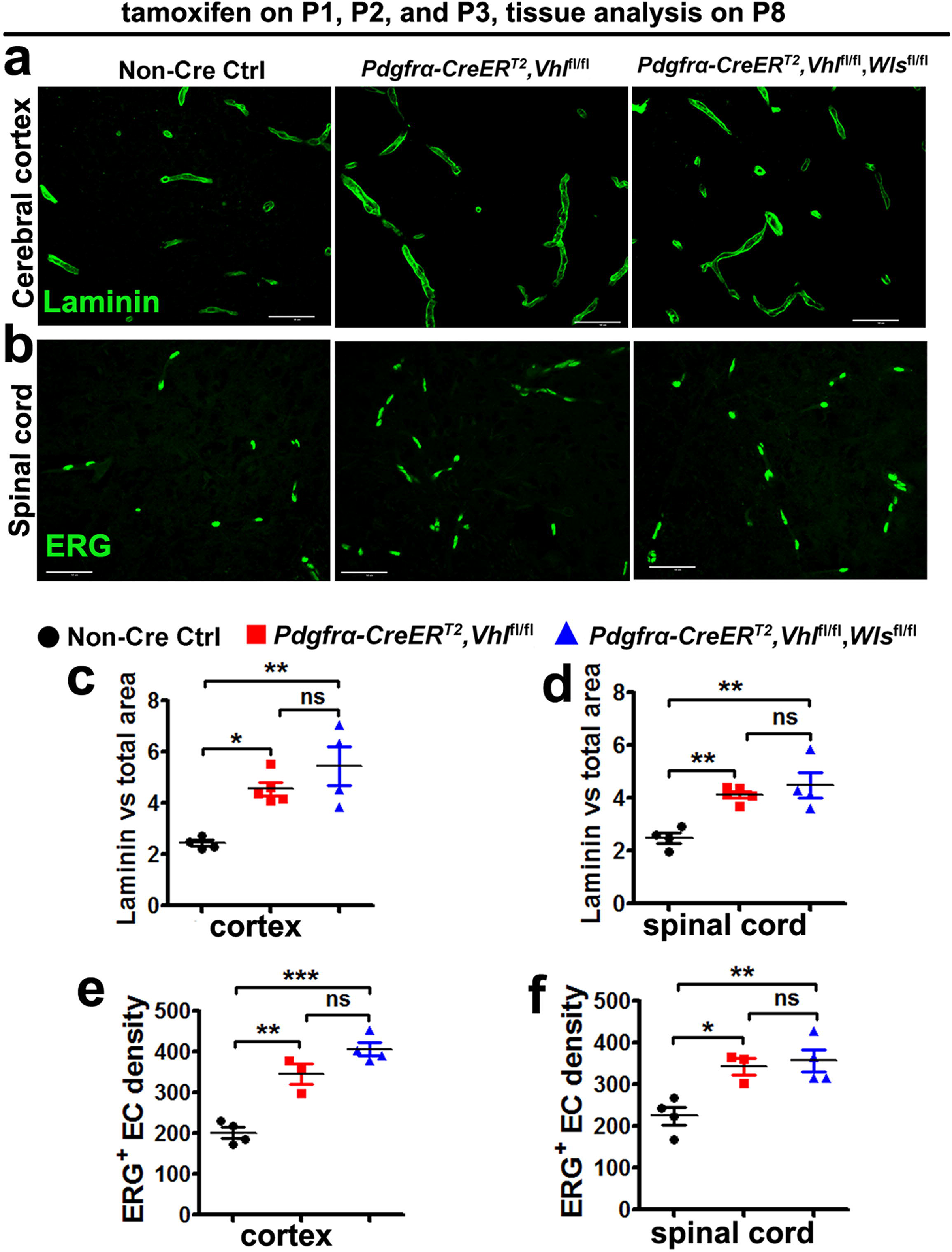
Blocking Wnt secretion from oligodendrocyte progenitor cells (OPCs) does not affect HIFα-regulated CNS angiogenesis. **a-b**, representative confocal images of Laminin-labeled blood vessels and ERG-labeled ECs in each group of mice. Scale bars = 50 µm. **c-d** percent of Laminin-occupying area among total assessed area. Welch’s ANOVA followed by unpaired *t* test with Welch’s correction, * P < 0.05, *** *P* < 0.001, ns, not significant. *W*_(2, 5.256)_ = 30.38, *P* =0.0013 for cortex. One-way ANOVA followed by Tukey’s multiple comparisons, ** *P* < 0.01, ns, not significant. *F*_(2,10)_ = 12.67, *P* = 0.0018 for spinal cord. **e-f**, densities (#/mm^2^) of ERG^+^ ECs. One-way ANOVA followed by Tukey’s multiple comparisons, * P < 0.05, ** *P* < 0.01, *** *P* < 0.001, ns, not significant. *F*_(2,8)_ = 39.80, *P* < 0.0001 cortex, *F*_(2,8)_ = 10.30, *P* = 0.0064 spinal cord. Mice from the above three groups were injected with tamoxifen at P1, P2, and P3 and sacrificed at P8.

### VEGFA expression is upregulated in the CNS of oligodendroglial HIFα-stabilized transgenic mice

A previous study reported that VEGFA was unperturbed by oligodendroglial HIFα stabilization ^9^. We revisited the potential connection between HIFα and VEGFA in oligodendrocytes both *in vivo* and *in vitro*. HIF1α cKO (or HIF2α cKO) alone did not alter *Vegfa* mRNA level in the CNS of *Cnp-Cre:Hif1α*^fl/fl^ (or *Cnp-Cre:Hif2α*^fl/fl^) mutants compared with non-Cre controls (data not shown), indicating a redundancy of oligodendroglial HIF1α and HIF2α in regulating VEGFA. HIFα double cKO (i.e. *Cnp-Cre:Hif1α*^fl/fl^:*Hif2α*^fl/fl^) decreased *Vegfa* mRNA expression in the spinal cord (Fig. 6a) and reduced the secretion of VEGFA from primary brain OPCs into the culture medium (Fig. 6b). Conversely, Genetic HIFα stabilization by VHL deletion increased *Vegfa* mRNA expression in the spinal cord of *Cnp-Cre:Vhl*^fl/fl^ animals (Fig. 6c, d1-d4). Double fluorescent in situ hybridization confirmed that *Vegfa* mRNA was upregulated in *Plp* mRNA^+^ oligodendroglial lineage cells *in vivo* (Fig. 6 e1-e2). Moreover, time-conditional and stage-specific VHL cKO demonstrated that genetic HIFα stabilization activated VEGFA not only in PDGFRα^+^ OPCs (*Pdgfrα-CreER^T2^:Vhl*^fl/fl^ strain) (Fig. 6f) but also in PLP^+^ oligodendroglia (*Plp-CreER^T2^:Vhl*^fl/fl^ strain) (Fig. 6g). Pharmacological DMOG treatment increased *Vegfa* mRNA expression and HIFα signaling blocker Chetomin prevented DMOG-induced *Vegfa* activation in primary OPCs purified from neonatal murine brains (Fig. 6h). Furthermore, *Vegfa* mRNA was elevated by greater than 3-fold in primary OPCs purified from neonatal *Sox10-Cre:Vhl*^fl/fl^ mice compared with those from non-Cre littermate controls (Fig. 6i). All these data suggest that VEGFA is regulated by HIFα in oligodendroglial lineage cells.

**Figure 6.**
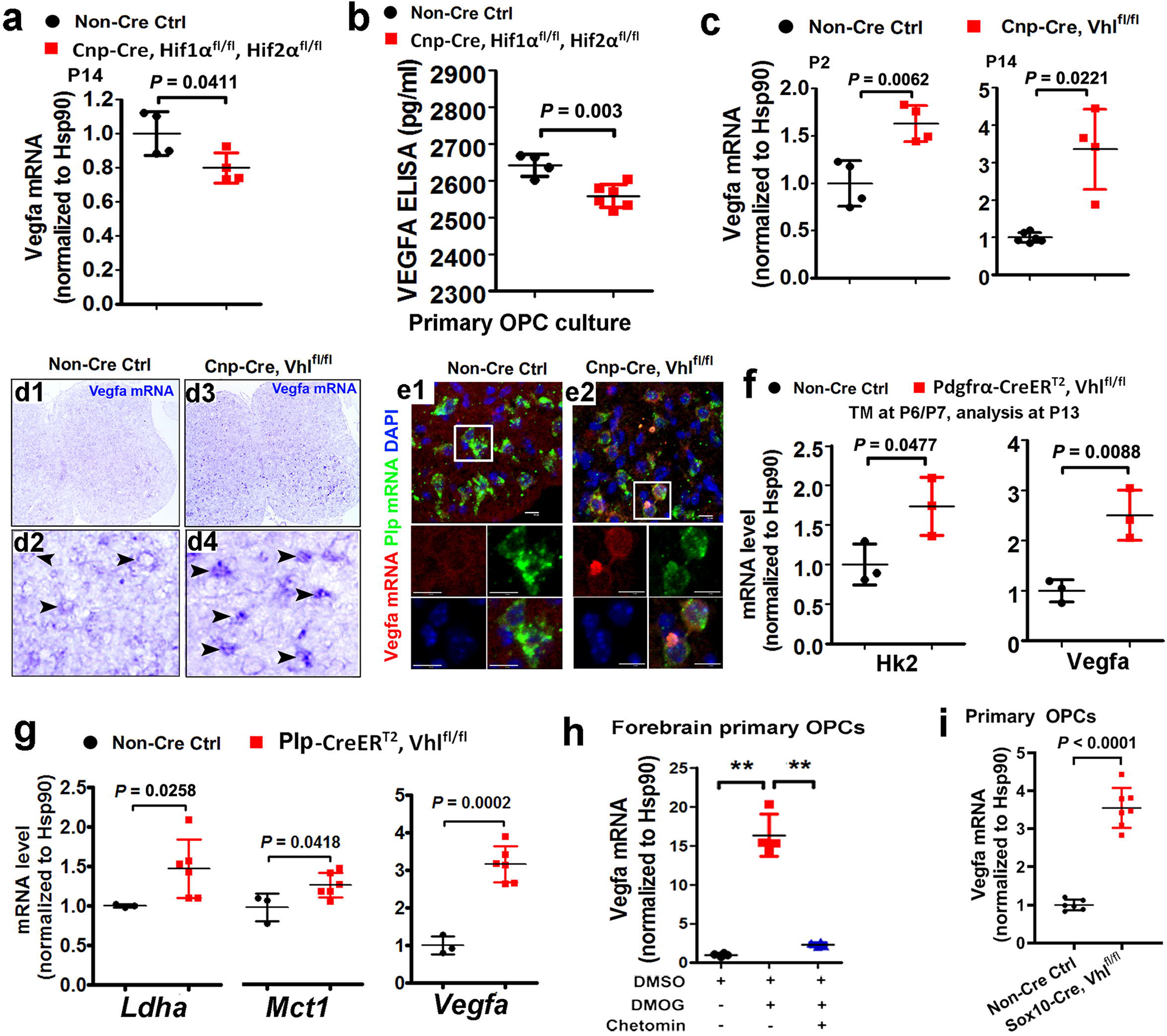
Oligodendroglial HIFα regulates VEGFA expression. **a**, RT-qPCR assay of *Vegfa* mRNA in the spinal cord of P14 mice. Two-tailed Student’s *t* test, t_(6)_ = 2.590. **b**, ELISA measurement of VEGFA concentration in the culture medium of primary OPCs isolated from neonatal forebrain of indicated genotypes. Two-tailed Student’s *t* test, t_(8)_ = 4.208. **c**, RT-qPCR assay of *Vegfa* mRNA in the spinal cord. Two-tailed Student’s *t* test, t_(6)_ = 4.127 at P2; Welch’s correction t_(3.139)_ = 4.299 at P14. **d1-d4**, *Vegfa* mRNA in situ hybridization in the spinal cord of P8 transgenic mice. Arrowheads point to *Vegfa* mRNA^+^ cells. Note that *Vegfa* mRNA signals were higher in the *Cnp-Cre, Vhl*^fl/fl^ spinal cord (d2) than those in the non-Cre controls (d4). **e1-e2**, *Vegfa* and *Plp* dual fluorescent mRNA in situ hybridization in P8 spinal cord. Boxed areas were shown at higher magnification in single color channels. Note that *Vegfa* mRNA signals are higher in *Plp*^+^ oligodendrocytes in *Cnp-Cre, Vhl*^fl/fl^ mutants than those in non-Cre controls. Scale bars: 10 µm. **f**, RT-qPCR assay of *Hk2* and *Vegfa* mRNA in the spinal cord of P14 *Pdgfrα-CreER*^T2^*, Vhl*^fl/fl^ and non-Cre littermate controls that had been treated with tamoxifen (TM) at P6 and P7. Two-tailed Student’s *t* test, t_(4)_ = 2.822 *Hk2*, t_(4)_ = 4.770 *Vegfa*. **g**, RT-qPCR assay of the mRNA levels of HIFα target genes *Ldha, and Mct1* and *Vegfa* in the spinal cord of P14 *Plp-CreER*^T2^*, Vhl*^fl/fl^ and non-Cre littermate controls that had been treated with tamoxifen at P6 and P7. Two-tailed Student’s *t* test, t_(7)_ = 2.487 *Mct1*, t_(7)_ = 7.194 *Vegfa*; Two-tailed Student’s *t* test with Welch’s correction, t_(5.079)_ = 3.116 *Ldha*. **h**, RT-qPCR assay of *Vegfa* mRNA in primary OPCs treated with HIFα stabilizer DMOG and inhibitor Chetomin (cf Supplementary Figure 9). Welch’s ANOVA followed by unpaired *t* test with Welch’s correction,** *P* < 0.01. *W*_(2, 5.270)_ = 79.93, *P* =0.0012. **i**, RT-qPCR assay of *Vegfa* mRNA in primary OPCs isolated from neonatal brains of Sox10-Cre, Vhl^fl/fl^ mutants and littermate controls. Two-tailed Student’s *t* test, Welch’s-corrected t_(7.008)_ = 12.38.

### Blocking oligodendroglia-derived VEGFA significantly reduces oligodendroglial HIFα-regulated CNS angiogenesis

The regulation of VEGFA by oligodendroglial HIFα led us to hypothesize that VEGFA may be a crucial downstream molecule that couples oligodendroglial HIFα and CNS endothelial cell proliferation and vessel formation. To test this hypothesis, we generated *Pdgfrα-CreER*^T2^:*Vhl*^fl/fl^:*Vegfa*^fl/fl^ (HIFα-stabilized/VEGFA-disrupted), *Pdgfrα-CreER*^T2^:*Vhl*^fl/fl^ (HIFα-stabilized), and non-Cre control mice (Fig. 7a, b). The mRNA level of EC-specific marker PECAM1 was significantly attenuated in the spinal cord of *Pdgfrα-CreER*^T2^:*Vhl*^fl/fl^: *Vegfa*^fl/fl^ mice compared with that of *Pdgfrα-CreER*^T2^:*Vhl*^fl/fl^ mice (Fig. 7c). The densities of blood vessels (Fig. 7d-e), ERG^+^ total ECs (Fig. 7f), and ERG^+^BrdU^+^ proliferating ECs (Fig. 7g) were all significantly reduced in the spinal cord and cerebral cortex of *Pdgfrα-CreER*^T2^:*Vhl*^fl/fl^:*Vegfa*^fl/fl^ mice compared with those of *Pdgfrα-CreER*^T2^:*Vhl*^fl/fl^ mice. These data demonstrate that VEGFA disruption attenuates oligodendroglial HIFα-regulated CNS angiogenesis, thus providing unambiguous *in vivo* data arguing for an essential role of VEGFA in coupling oligodendroglial HIFα function and CNS angiogenesis.

**Figure 7.**
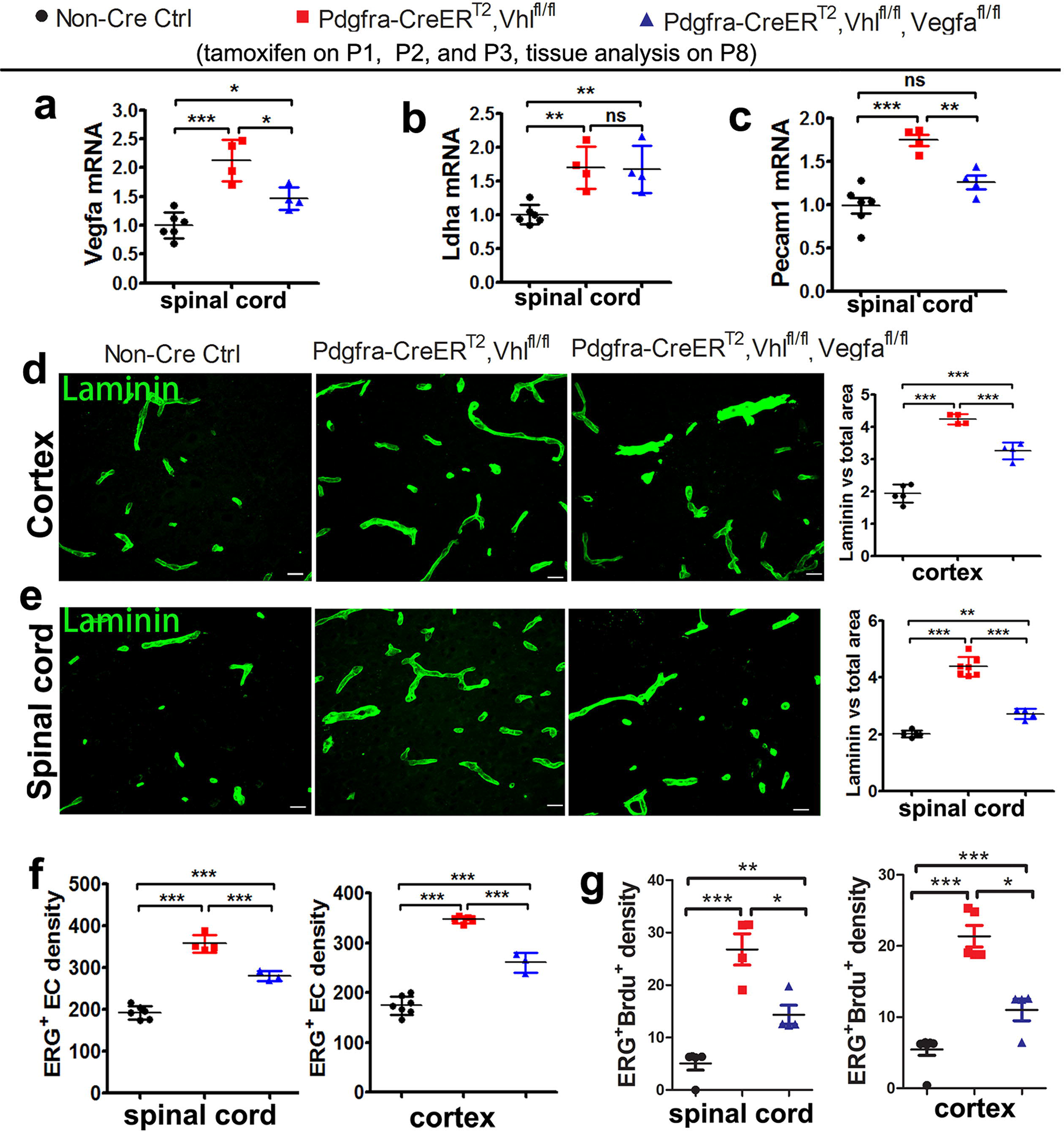
Oligodendroglial HIFα regulates CNS angiogenesis through VEGFA-mediated signaling. **a-c**,RT-qPCR assay of the mRNA levels of *Vegfa, Ldha, and Pecam1* in the spinal cord of each group of mice. One-way ANOVA followed by Tukey’s multiple comparisons, * *P* < 0.05, ** *P* < 0.01, ***, *P* < 0.001, ns not significant. *F*_(2, 11)_ = 21.74, *P* = 0.0002 *Vegfa; F*_(2, 11)_ = 11.50, *P* = 0.0020 *Ldha*; *F*_(2, 11)_ = 20.31, *P* = 0.0002 *Pecam1*. **d-e,** representative confocal images and quantification of Laminin-positive blood vessels. One-way ANOVA followed by Tukey’s multiple comparisons, ** *P* < 0.01, ***, *P* < 0.001, ns not significant. *F*_(2, 10)_ = 105.5, *P* < 0.0001 cortex; *F*_(2, 13)_ = 133.2, *P* < 0.0001 spinal cord. Scale bars = 10 μm. **f-g**, densities (#/mm^2^) of ERG^+^ ECs and ERG^+^BrdU^+^ proliferating ECs. One-way ANOVA followed by Tukey’s multiple comparisons, * *P* < 0.05, ** *P* < 0.01, ***, *P* < 0.001. ERG^+^ ECs, *F*_(2, 10)_ = 113.1, *P* < 0.0001 spinal cord, *F*_(2, 12)_ = 169.0, *P* < 0.0001 cortex; ERG^+^BrdU^+^ proliferating ECs, *F*_(2, 10)_ = 24.09, *P* = 0.0001 spinal cord, *F*_(2, 12)_ = 86.14, *P* < 0.0001 cortex.

### Blocking Wnt secretion from astroglia attenuates astroglial HIFα-regulated CNS angiogenesis

Astroglial maturation is also temporally and functionally coupled with postnatal CNS angiogenesis. We assess the connection of astroglial HIFα and Wnt/β-catenin activation in the CNS. We first used the mouse *Gfap* promoter-driven constitutive Cre, i.e. *mGfap-Cre* to genetically stabilize HIFα in astroglia. The efficiency of *mGfap-Cre*-mediated recombination among astroglial lineage cells, quantified by Cre-mediated EYFP reporter, was low (~35%) in the CNS in the early postnatal CNS by P10 (Supplementary Fig. 12a-c) and progressively increased during postnatal CNS development (supplementary Fig. 12d-f). Our fate-mapping data showed that EYFP reporter, which is an indicator of *mGfap-Cre* activity, was expressed in GFAP^+^ or S100β^+^ astrocytes, but not in Sox10^+^ oligodendroglial lineage cells, NeuN^+^ neurons (Fig. 8a), or ERG^+^ ECs (data not shown) in the spinal cord and the cerebral cortex of adult *mGfap-Cre:Rosa26-EYFP* mice at P60, confirming that *mGfap-Cre* primarily targets astroglial lineage cells in those CNS regions.

**Figure 8.**
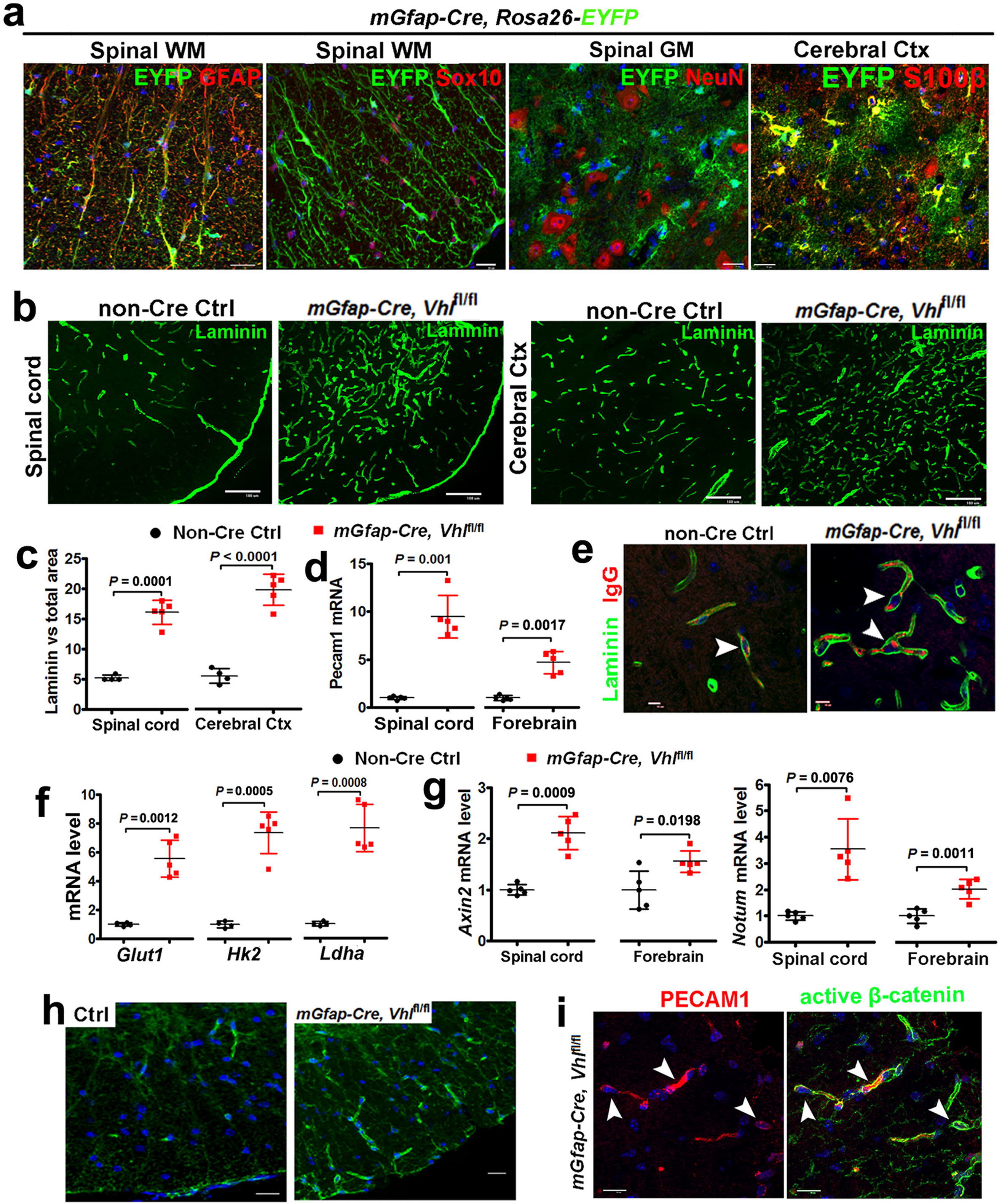
Astroglial HIFα stabilization promotes CNS angiogenesis and enhances Wnt signaling activity. **a**, fate-mapping study showing that mGfap-Cre-mediated EYFP was expressed in GFAP^+^ astrocytes but not in Sox10^+^ oligodendroglial lineage cells or NeuN^+^ neurons in the spinal cord at P60. WM, white matter, GM, gray matter, Ctx, cortex. EYFP was identified as S100β^+^ astrocytes in the cerebral Ctx. Scale bars = 20 μm. **b**, representative images of Laminin immunostaining in *mGfap-Cre, Vhl*^fl/fl^ mutants and non-Cre control mice at P30. Scale bars = 100 μm. **c**, percentage of Laminin-occupying area among total area at P30. Two-tailed Student’s *t* test, Welch’s corrected *t*_(4)_ = 11.92 spinal cord, *t*_(7)_ = 10.12 cerebral cortex. **d**, RT-qPCR assay of endothelial Pecam1 at P30. Two-tailed Student’s *t* test, Welch’s corrected *t*_(4.046)_ = 8.564 spinal cord, Welch’s corrected *t*_(4.490)_ = 6.706 forebrain. **e**, immunostaining showing that endogenous mouse IgG is restricted to Laminin^+^ blood vessels (arrowheads) in the early adult spinal cord of *mGfap-Cre,Vhl*^fl/fl^ mutant and control mice at P47. Scale bars = 10 μm. **f**, RT-qPCR assay of the mRNA levels of HIFα target gene *Glut1, Hk2*, and *Ldha* in P30 spinal cord. Two-tailed Student’s *t* test with Welch’s correction, *t*_(4.099)_ = 7.947 *Glut1*, *t*_(4.243)_ = 9.636 *Hk2*, *t*_(4.085)_ = 9.025 *Ldha.* **g**, RT-qPCR assay of the mRNA levels of Wnt/β-catenin target genes *Axin2* and *Notum* at P30. Two-tailed Student’s *t* test, Welch’s corrected *t*_(4.760)_ = 7.296 spinal cord *Axin2*, *t*_(8)_ = 2.902 forebrain *Axin2*, Welch’s corrected *t*_(4.149)_ = 4.850 spinal cord *Notum*, t_(8)_ = 4.994 forebrain *Notum*. **h**, immunostaining of active β-catenin in the spinal cord of mGfap-Cre, Vhl^fl/fl^ mutants and non-Cre control mice at P30. Scale bars = 10 μm. **i**, double immunostaining of active β-catenin and PECAM1 in the spinal cord of *mGfap-Cre, Vhl*^fl/fl^ mutants at P30. Arrowheads point to double positive cells. Blue is DAPI nuclear staining. Scale bars = 10 μm.

We observed a significant increase in the density of Laminin^+^ blood vessels (Fig. 8b, c) and in the mRNA expression of EC-specific PECAM1 (Fig. 8d) throughout the CNS of *mGfap-Cre:Vhl*^fl/fl^ mutants compared with non-Cre control mice by P30 when Cre-mediated recombination efficiency was greater than 80% (Fig. 8a, Supplementary Fig. 12). Double immunohistochemistry showed that blood-borne macromolecule IgG was confined to Laminin^+^ blood vessels in *mGfap-Cre:Vhl*^fl/fl^ mice, a similar pattern to that in age-matched non-Cre controls (Fig. 8e, arrowheads), indicating that the function of the blood brain (spinal cord) barrier does not appear compromised although the vessel density is elevated.

Unexpectedly, we found that stabilizing HIFα in astroglial lineage cells (Fig. 8f) remarkably activated Wnt/β-catenin signaling in the CNS of *mGfap-Cre:Vhl*^fl/fl^ mice, as shown by significant elevation in the expression of Wnt/β-catenin signaling target genes *Axin2* and *Notum* in spinal cord and brain (Fig. 8g). Histological (Fig. 8h) and Western blot (cf Fig. 9a, b) assay demonstrated that the active form of β-catenin (dephosphorylated on Ser37 or Thr41) was significantly increased in *mGfap-Cre:Vhl*^fl/fl^ mice. Double immunohistochemistry confirmed the presence of elevated active β-catenin in PECAM1^+^ ECs (Fig. 8i, arrowheads). Collectively, our data suggest that stabilizing HIFα in astroglial lineage cells increases CNS angiogenesis and activates Wnt/β-catenin signaling in ECs.

**Figure 9.**
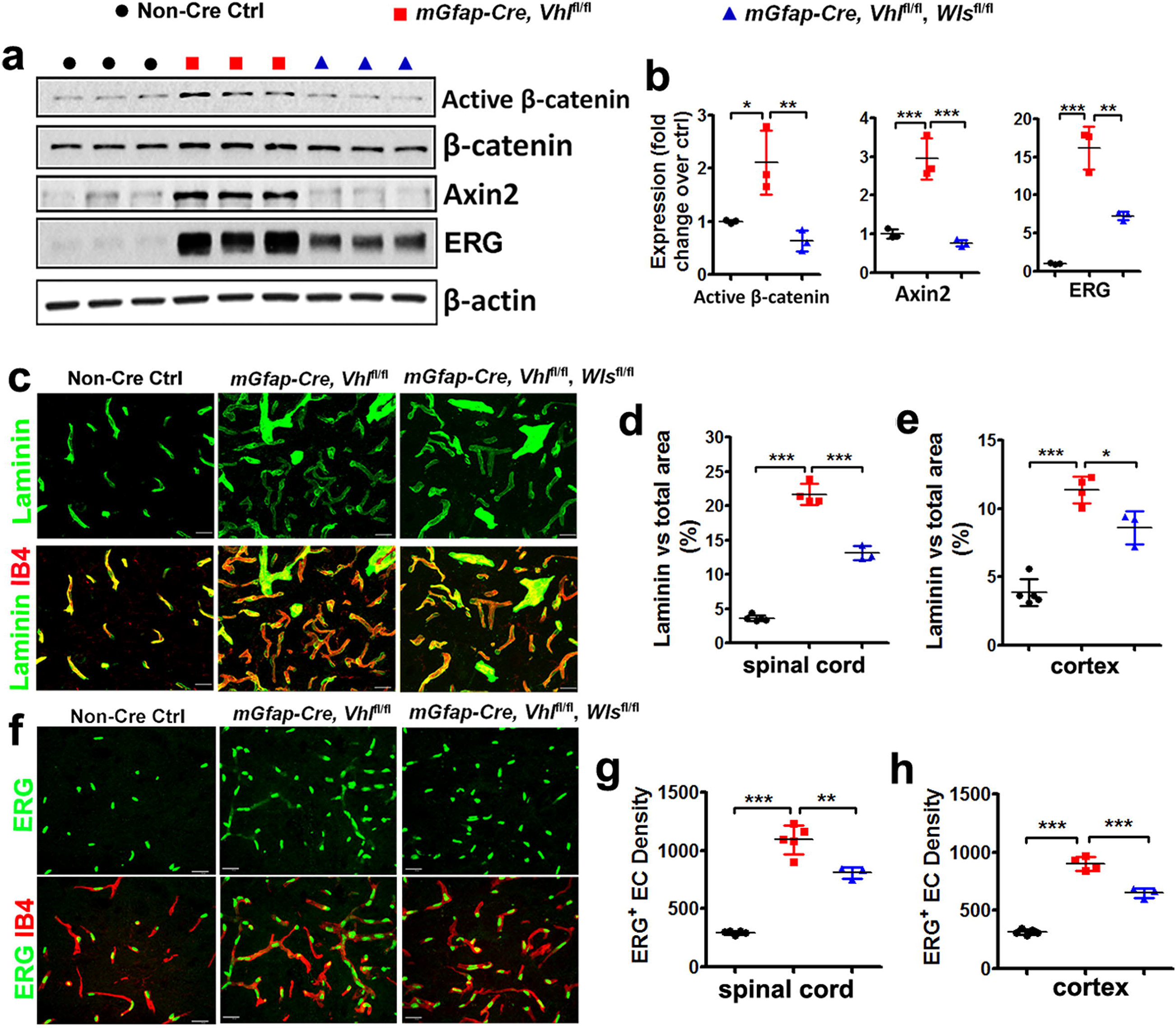
Constitutively blocking Wnt secretion from astrocytes reduces HIFα-regulated angiogenesis in the early adult CNS. **a-b**, Western blot (**a**) and quantification (**b**) of the active form of β-catenin, Wnt/β-catenin target gene Axin2, EC-specific nuclear protein ERG in the spinal cord at P30. One-way ANOVA followed by Tukey’s multiple comparisons, * *P* < 0.05, ** *P* < 0.01, *** *P* < 0.001. *F*_(2, 6)_ = 13.38, *P* = 0.0061 active β-catenin, *F*_(2, 6)_ = 41.26, *P* = 0.0003 Axin2, *F*_(2, 6)_ = 65.59, *P* < 0.0001 ERG, **c**, representative confocal images of Laminin and IB4 in the spinal cord at P30, scale bars = 20 μm. **d-e**, percentage of Laminin-occupying area among total area at P30. One-way ANOVA followed by Tukey’s multiple comparisons, * *P* < 0.05, *** *P* < 0.001. *F*_(2, 9)_ = 332.0, *P* < 0.0001 spinal cord, *F*_(2, 9)_ = 59.19, *P* < 0.0001 cerebral cortex. **f**, representative confocal images of ERG and IB4 in the spinal cord at P30, scale bars = 20 μm. **g-h**, densities (per mm^2^) of ERG^+^ ECs at P30. One-way ANOVA followed by Tukey’s multiple comparisons, ** *P* < 0.01, *** *P* < 0.001. *F*_(2, 10)_ = 124.1, *P* < 0.0001 spinal cord, *F*_(2, 9)_ = 219.2, *P* < 0.0001 cerebral cortex.

Wnt/β-catenin signaling activation in ECs by astroglial HIFα stabilization led us to hypothesize that astroglia-derived Wnt signaling may instead play a major role in HIFα-regulated CNS angiogenesis. To test this hypothesis, we generated *mGfap-Cre:Vhl*^fl/fl^:*Wls*^fl/fl^ mice to stabilize HIFα’s function and simultaneously disrupting Wnt secretion from HIFα-stabilized astroglia. Our data showed that Wnt signaling activity was significantly reduced in the spinal cord of *mGfap-Cre:Vhl*^fl/fl^:*Wls*^fl/fl^ mice compared with that of *mGfap-Cre:Vhl*^fl/fl^ mice (Fig. 9a, b), thus verifying the efficacy of blocking astroglia-derived Wnt signaling *in vivo* by WLS deletion. Intriguingly, disrupting astroglia-derived Wnt signaling significantly reduced the densities of blood vessels (Fig. 9c-e) and ERG^+^ ECs (Fig. 9f-h) in the CNS of *mGfap-Cre:Vhl*^fl/fl^:*Wls*^fl/fl^ double mutant mice compared with *mGfap-Cre:Vhl*^fl/fl^ mice, indicating that astroglia-derived Wnt signaling is a downstream mediator of astroglial HIFα-regulated CNS angiogenesis.

Our results indicated that the constitutive mGfap-Cre elicited a poor recombination efficiency in early postnatal astrocytes (Supplementary Fig. 12). To determine whether early postnatal astrocytes regulate CNS angiogenesis through HIFα-activated Wnt signaling, we generated *Aldh1l1-CreER^T2^*:*Vhl*^fl/fl^:*Wls*^fl/fl^ mutants. Our data demonstrated a greater than 90% of recombination efficiency and 95% of astroglial specificity in the spinal cord and cerebral cortex of *Aldh1l1-CreER^T2^*:Rosa26-EYFP at P8 when tamoxifen was injected at P1, P2, and P3 (Supplementary Fig. 13). Consistent with the data derived from *mGfap-Cre:Vhl*^fl/fl^:*Wls*^fl/fl^ strain, we found that the densities of blood vessels and ECs were significantly increased in *Aldh1l1-CreER^T2^*:*Vhl*^fl/fl^ mutants compared with those in non-Cre controls and that simultaneous WLS ablation significantly reduced the densities of blood vessels and ECs in the cortex and spinal cord of *Aldh1l1-CreER^T2^*:*Vhl*^fl/fl^:*Wls*^fl/fl^ mutants compared with those of *Aldh1l1-CreER^T2^*:*Vhl*^fl/fl^ animals at early postnatal age of P8 (Fig. 10). These data provide a strong genetic proof that HIFα-activated Wnt signaling is a major downstream pathway by which astroglia regulate angiogenesis during postnatal CNS development.

**Figure 10.**
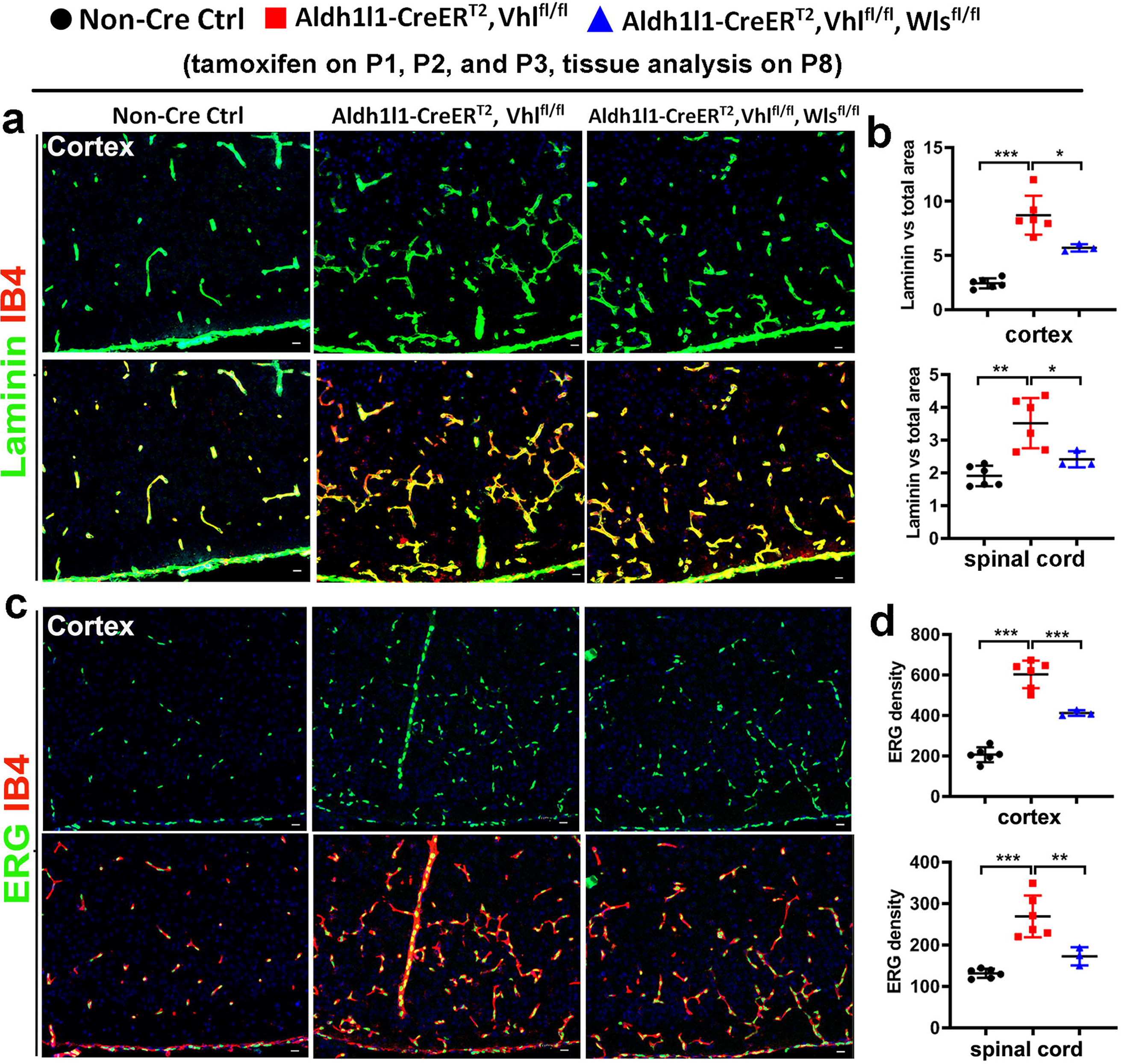
Conditionally blocking Wnt secretion from neonatal astrocytes reduces HIFα-regulated CNS angiogenesis in the early postnatal CNS. **a,** representative confocal images of Laminin and IB4 in the cerebral cortex of each group of mice at P8 that had been treated with tamoxifen at P1, P2, and P3. **b**, percentage of Laminin-occupying area among total area at P8. Cortex, one-way ANOVA followed by Tukey’s multiple comparisons, * *P* < 0.05, *** *P* < 0.001, *F*_(2, 12)_ = 41.00, *P* < 0.0001. Spinal cord, Welch’s ANOVA followed by unpaired t test with Welch’s correction, * *P* < 0.05, ** *P* < 0.01 *W*_(2, 6.824)_ = 11.42, *P* = 0.0067. **c**, representative confocal images of Laminin and IB4 in the P8 cerebral cortex of each group of mice that had been treated with tamoxifen at P1, P2, and P3. **d**, density (#/mm^2^) of ERG^+^ ECs at P8. One-way ANOVA followed by Tukey’s multiple comparisons, ** *P* < 0.01, *** *P* < 0.001. *F*_(2, 12)_ = 92.24, *P* < 0.0001 cortex, *F*_(2, 12)_ = 24.37, *P* < 0.0001 spinal cord.

## Discussion

The maturation of glial cells including oligodendroglia and astroglia in the developing human and murine brain is temporally and functionally coupled with the maturation of the CNS vascular network^21^. The regulation of CNS angiogenesis by glial cells is critical for postnatal CNS development and investigating the molecular underpinnings of CNS angiogenesis has clinical implications in neural repair after CNS damage in which hypoxia is commonly present ^4,21,22^. In this study, we employed a battery of genetic mutant mice and presented several significant and novel findings: 1) oligodendroglial HIFα is necessary and sufficient for postnatal CNS angiogenesis and this regulation occurs in a manner independent of CNS regions; 2) In sharp contrast to previous report^9^, HIFα stabilization in oligodendroglial lineage cells does not perturb Wnt/β-catenin signaling, but remarkably activates VEGF, and genetically blocking oligodendroglia-derived VEGF but not Wnt reduces oligodendroglial HIFα-regulated CNS angiogenesis; 3) Wnt signaling is a downstream pathway by which astroglial HIFα regulates CNS angiogenesis. Our findings represent an alternative view in our mechanistic understanding of oligodendroglial HIFα-regulated angiogenesis from a Wnt-dependent/VEGF-independent view^9^ to a VEGF-dependent/Wnt-independent one, and also unveil a glial cell type-dependent HIFα-Wnt axis (oligodendroglial vs astroglia) in regulating CNS angiogenesis (Supplementary Fig. 14).

Previous data suggest that the regulation of Wnt/β-catenin signaling (activation or repression) by HIFα is cell type and/or context-dependent^10,23–27^. It is important to determine whether HIFα in CNS glial cells differentially regulates Wnt/β-catenin signaling *in vivo*. A recent study reported that oligodendroglial HIFα could activate Wnt/β-catenin signaling not only in oligodendroglial lineage cells but also in endothelial cells through HIFα-mediated Wnt7a/7b expression^9^. However, the *in vivo* and *in vitro* data presented in this study do not support this assertion. In our study, we employed five different strains of oligodendroglial Cre (two constitutive Cre and three inducible Cre) to genetically stabilize HIFα and found no evidence of Wnt/β-catenin activation or Wnt7a/7b upregulation in the brain and the spinal cord and in primary OPCs. The failure to detect Wnt/β-catenin activation is unlikely due to the inefficiency of HIFα stabilization because the canonical HIFα target genes, for example, those which are involved in glycolysis, are consistently and significantly upregulated in our transgenic animals and in primary OPCs, thus verifying the efficacy of HIFα stabilization. In sharp contrast, we found that Wnt/β-catenin signaling activity is significantly upregulated in the CNS of astroglial HIFα-stabilized mice, suggesting that our experimental approach is effective in quantifying the changes of Wnt/β-catenin activity and that the regulation of Wnt/β-catenin signaling by HIFα is glial cell type-dependent in the CNS.

Inspired by glial cell type (oligodendroglia vs astroglia)-dependent activation of Wnt/β-catenin signaling, we proposed a working model in which HIFα-activated Wnt signaling regulates endothelial cell proliferation and vessel formation in a cell type-dependent manner (Supplementary Fig. 14). To avoid the intrinsic caveats of pharmacological compounds and *in vitro* culture systems, we employed *in vivo* genetic models of VHL/WLS double cKO to stabilize HIFα and simultaneously disrupt the secretion of Wnt ligands. Indeed, our data demonstrate that WLS-deficiency decreases Wnt secretion and Wnt7a-induced autocrine Wnt/β-catenin signaling in primary OPCs and that disrupting WLS in astroglia reduces the activity of astroglial HIFα-regulated Wnt/β-catenin signaling in the CNS. Based on VHL/WLS double cKO systems, we provide strong genetic evidences that HIFα-regulated Wnt signaling from astroglia but not oligodendroglia plays a crucial role in regulating postnatal CNS angiogenesis. Our findings argue against a major role oligodendroglia-derived/HIFα-activated Wnt/β-catenin signaling in angiogenesis in the developing murine CNS as previously reported ^9^.

There are 19 Wnt ligands in rodents, which can be grossly classified into canonical and non-canonical sub-types depending on the necessity of β-catenin for the signaling activation^10^. WLS ablation blocks the secretion of all Wnt members from Wnt-producing cells^16–19,28^. In our genetic models, we demonstrate that ablation of WLS remarkably reduces astroglial HIFα-mediated canonical Wnt/β-catenin signaling. However, we cannot exclude the possibility that astroglia-derived non-canonical Wnt signaling is also altered in our genetic manipulation which may be potentially involved in CNS angiogenenic regulation^29^. Moreover, it remains unclear which Wnt ligand(s) plays a major role in coupling astroglial HIFα with CNS angiogenesis. Our preliminary data do not support a major role of Wnt7a/7b, because neither Wnt7a nor Wnt7b expression was activated in oligodendroglial and astroglial HIFα-stabilized CNS. Future studies are needed to pinpoint which Wnt ligand(s) are the downstream mediator(s) of astroglial HIFα-regulated CNS angiogenesis.

VEGF (i.e. VEGFA) is a well-established angiogenic and neurotrophic factor in the CNS^4,30–35^. VEGFA regulates angiogenesis in the developing and adult CNS through its membrane-bound receptors VEGFR-1(Flt1) and VEGFR-2 (Kdr)^30^. In the early postnatal CNS, VEGFR-1 and 2 are highly expressed in the vascular ECs. However, the ligand VEGFA is barely detectable in the vascular ECs but highly expressed in the parenchymal neural cells including oligodendroglial lineage cells^20,36–38^. Previous study suggested that the HIFα-VEGF connection did not occur in the CNS oligodendroglia^9^ although this connection was demonstrated in the retina^39^. In this study, we found that oligodendroglial HIFα cKO reduces VEGFA whereas oligodendroglial HIFα stabilization increases VEGFA expression, indicating that HIFα transcriptionally regulates VEGFA in oligodendroglial lineage cells. Further corroborating these findings, purified primary OPCs respond to HIFα signaling stabilizer (DMOG) and blocker (Chetomin) by activating and inactivating VEGFA, respectively. Our results are consistent with previous data showing that VEGFA is a direct transcriptional target of HIFα^40–42^. By leveraging our unique *in vivo* genetic models of VHL/VEGFA double cKO, we unequivocally prove that VEGFA is an essential downstream molecule that couples oligodendroglial HIFα function and vascular angiogenesis in the CNS, which is different from Yuen et al.,^9^ who reported that VEGFA was unchanged in the CNS of oligodendroglial HIFα-stabilized mutants. The discrepancy may presumably reflect the intrinsic differences of *in vitro* pharmacological interventions and *in vivo* genetic manipulations.

It has been suggested that Wnt signaling regulates VEGF, or vice versa, to control angiogenesis^43–45^. It is possible that Wnt/β-catenin signaling is required for, or synergistically regulates, HIFα-activated VEGFA expression. Our data do not support this possibility. First, stabilizing oligodendroglial HIFα activates VEGFA but not Wnt/β-catenin signaling. Second, VEGF expression is indistinguishable in the CNS of oligodendroglial VHL/WLS double cKO mutants from that of oligodendroglial VHL single cKO mutants (data not shown), indicating that oligodendroglial-derived Wnt signaling plays a minor role in VEGFA expression. Third, Wnt/β-catenin activity is comparable in the CNS of oligodendroglial VHL/VEGFA double cKO mutants and VHL single cKO mutants, implying that oligodendroglia-derived VEGFA has no regulatory role in Wnt/β-catenin activity. Together, our results does not support a major interplay between oligodendroglial HIFα-activated VEGFA and Wnt/β-catenin signaling in modulating CNS angiogenesis.

Previous studies including those from our own laboratory^10,12,46–49^ have shown that dysregulated Wnt/β-catenin activity invariably inhibits oligodendrocyte differentiation and myelination. Given the normal level of Wnt/β-catenin activity in the CNS of oligodendroglial HIFα-stabilized mutants, our study will spark renewed interests in studying Wnt-independent mechanisms underlying the impairment of oligodendroglial differentiation and myelination in HIFα-stabilized mutants as previously reported^9^. Interestingly, HIFα is stabilized and enriched in oligodendroglia in the active demyelinating lesions and normal appearing white matter (NAWM) of multiple sclerosis patient brains^50–53^. We show that HIFα stabilization in oligodendrocytes remarkably activates the angiogenic and neurotrophic factor VEGF in the CNS. The genetic models generated in our study also provide a powerful tool in determining the role of HIFα stabilization in OPCs and oligodendrocytes in the pathophysiology of demyelination and remyelination in multiple sclerosis and other neurological disorders in which hypoxia-like tissue injury occurs.

## Methods

### Animals

A total of 14 transgenic strains were used in this study. Cnp-Cre mice (RRID: MGI_3051754)^11^, Sox10-Cre (RRID: IMSR_JAX:025807), Sox10-CreERT2 (RRID:IMSR_JAX: 027651), Pdgfrα-CreERT2 (RRID: IMSR_JAX:018280), Aldh1l1-CreERT2 (RRID: IMSR_JAX:029655), Plp-CreERT2 (RRID: IMSR_JAX:005975), mGfap-Cre (RRID: IMSR_JAX:024098), Hif1α-floxed (RRID: IMSR_JAX:007561), Hif2α-floxed (RRID: IMSR_JAX:008407), Vhl-floxed (RRID: IMSR_JAX:012933), Wls-floxed (RRID: IMSR_JAX:012888), Vegfa-floxed (MGI:1931048)^54^, Bat-lacZ (RRID: IMSR_JAX:005317), Rosa26-EYFP (RRID: IMSR_JAX:006148). Animals were housed at 12h light/dark cycle with free access to food and drink, and both males and females were used in this study. The single transgenic mice were crossed to generate double or triple transgenic mice indicated in the study. All Cre transgene was maintained as heterozygous. All transgenic mice were maintained on a C57BL/6 background. Animal protocols were approved by Institutional Animal Care and Use Committee at the University of California, Davis.

Primers used for genotyping *Hif1α*-floxed, *Hif2α*-floxed and *Vhl*-floxed and for detecting Cre-mediated DNA deletion (as in Fig. 1A and Fig. 2A-B) were derived from previous study ^55^ and listed here. *Hif1α*-F1: 5′-TTGGGGATGAAAACATCTGC-3′, *Hif1α*-F2: 5′-GCAGTTAAGAGCACTAGTTG-3′, *Hif1α*-R: 5′-GGAGCTATCTCTCTAGACC-3’), *Hif2α*-F1:5′-CAGGCAGTATGCCTGGCTAATTCCAGTT-3′, *Hif2α*-F2: 5′-CTTCTTCCATCATCTGGGATCTGGGACT-3′, *Hif2α*-R: 5′-GCTAACACTGTACTGTCTGAAAGAGTAGC-3′, *Vhl*-F1: 5′-CTGGTACCCACGAAACTGTC-3′, *Vhl*-F2: 5′-CTAGGCACCGAGCTTAGAGGTTTGCG-3′ *Vhl*-R: 5′-CTGACTTCCACTGATGCTTGTCACAG-3′. Primers used for genotyping Wls-floxed for detecting Cre-mediated DNA deletion (as in Fig. 7B) were derived from previous study ^18^ and listed here. *Wls*-F1: 5’-CTTCCCTGCTTCTTTAAGCGTC-3’, *Wls*-F2: 5’-AGGCTTCGAACGTAACTGACC-3’, *Wls*-R: 5’-CTCAGAACTCCCTTCTTGAAGC-3’

### Tamoxifen and BrdU (or EdU) treatment

Tamoxifen (TM) (T5648, Sigma) was dissolved in mixture of ethanol and sunflower oil (1:9 by volume) at a concentration of 30 mg/ml ^14^. Mice were administrated intraperitoneally (i.p.) with tamoxifen at dose of 200 μg/g body weight at time points indicated in each figure. BrdU (B5002, Sigma) or EdU (A10044, Thermo Fisher Scientific) was freshly dissolved in 0.9% sterile saline at a concentration of 10 mg/ml. BrdU or EdU was i.p. injected to animals at a dose of 100 ug/g body weight at time-points indicated in the figures. Tamoxifen was administered to both the controls and the inducible floxed mice in all experiments involving inducible Cre-LoxP approach.

### Primary OPC culture

Primary mixed glial culture (MG) was prepared from the forebrains of neonatal pups between ages P0 and P2. The isolated cortical tissues were dissociated by papain dissociation kit (#LK003176, Worthington) supplemented with DNase I (250U/ml; #D5025, Sigma) and D-(+)-glucose (0.36%; #0188 AMRESCO) in 33 ºC/10% CO_2_ for 90 min. Next, the tissues were transferred in PDS Kit-Inhibitor solution (#LK003182, Worthington). Tissue chunks were triturated, and then collect the cell suspension supernatant. After centrifugation, cells were plated on poly-D-lysine (PDL, #A003-E, Millipore)-coated 10 cm dishes (#130182, Thermo Scientific) in high glucose DMEM medium (#1196092, Thermo Fisher) with 10% heat-inactivated fetal bovine serum (#12306-C, Sigma) and penicillin/streptomycin (P/S, #15140122, Thermo Fisher). After 24h, attached cells were washed with HBSS (#24020117, Thermo Fisher) to remove serum, and maintained with serum free growth medium (GM), a 3:7 mixture (v/v) of B104 neuroblastoma-conditioned medium, 10 ng/ml Biotin (#B4639, Sigma), and N1 medium (high glucose DMEM supplemented with 5 μg/ml insulin (#I6634, Sigma), 50 μg/ml apo-transferrin (#T2036, Sigma), 100 μM Putrescine (#P5780, Sigma) 30 nM Sodium selenite (#S5261, Sigma), 20 nM Progesterone (#P0130, Sigma). We performed immunopanning 96 h after GM maintenance. Before immunopanning, cells were resuspended in panning solution (0.1% BSA in N1 medium). The cells were panned once with the anti-Thy1.2 antibody (#105302, Biolegend) for negative immunopanning and then panned once with the anti-NG2 antibody (#AB5320, Millipore) for positive immunopanning. OPCs were cultured on PDL-coated plates with complete GM. The complete GM consisted of GM with 5 ng/ml FGF(#450-33, Peprotech), 4 ng/ml PDGF-AA (#315-17, Peprotech), 50 µM Forskolin (#6652995, Peprotech,) and glutamax (#35050, Thermo Fisher). To induce differentiation, the medium was switched to differentiation medium (DM), which consists of 12.5 μg/ml insulin, 100 μM Putrescine, 24 nM Sodium selenite, 10 nM Progesterone, 10 ng/ml Biotin, 50 μg/ml Transferrin (#T8158, Sigma), 30 ng/ml 3,3’,5-Triiodo-L-thyronine (#T5516, Sigma), 40 ng/ml L-Thyroxine (#T0397, Sigma-Aldrich), glutamax and P/S in F12/high-glucose DMEM, 1:1 in medium (#11330032, Thermo Fisher Scientific).

### HIFα signaling stabilization and blockade *in vitro* by pharmacological approaches

Purified brain primary OPCs were pre-incubated with 100nM Chetomin or DMSO control for 2h, and then switched to the fresh culture medium with 1mM Dimethyloxalylglycine (DMOG, D3695, Sigma) in the presence of 100nM Chetomin (C9623, Sigma) or DMSO (D8418, Sigma) control for 7 hours before RNA preparation.

### VEGFA ELISA assay of primary OPCs

Cell culture medium of primary OPCs from Non-Cre control and *Cnp-Cre, Hif1α*^fl/fl^, were collected for VEGF measurement. Endogenous VEGF concentrations were determined using a mouse-specific VEGF Quantikine ELISA kit (t#MMV00, R&D System) according to the manufacture’s instruction.

### Transfection of Wnt7a and Wls-shRNA in primary OPCs and ELISA measurement of Wnt7a in culture medium

Primary OPCs was transfected with Wls-ShRNA (TRCN 0000 234932, Mission ShRNA bacterial Glycerol stock NM_026582) and ShRNA scramble control (Mission TRC2 PlkO.5-PURO Non-Mammalian shRNA control Plasmid), Wnt7a plasmid pLCN-Wnt7a-HA (Addgene, #18036) and empty pLCN-exp (Addgene, #64865) at the time points indicated in Figure 3a. The transfection was done by using FuGENE6 Transfection reagent (Promega, #E2691, lot#000371257). The Wnt7a in OPCs cell medium was measured by using mouse Wnt7a ELISA kit (Cusabio, #CSB-EL026141MO, Lot#G19147708) according to the manual of the kit.

### Tissue processing, immunohistochemistry, and semi-automated quantification of blood vessel density

Study mice were perfused with ice-cold phosphate buffer saline (PBS, PH7.0, Catalog #BP399-20, Fisher Chemical), and then post-fix in fresh 4% paraformaldehyde (PFA, Catalog #1570-S, Electron Microscopy Science, PA) at room temperature(RT) for 2 hours. The CNS tissue was washed in ice-cold PBS for three times, 15 minutes each time. The samples were cryoprotected with 30% sucrose in PBS (Sucrose, Catalog #S5-3, Fisher Chemical) for 20 hours followed by sectioning. Sixteen micron thick sections were serially collected and stored in −80 °C. Immunohistochemistry was conducted according to our previous studies ^56,57^. The information of primary antibodies used for immunohistochemistry in the study were listed in Supplementary Table 1. For BrdU immunostaining, sections were pretreated with fresh made 2N HCl (#320331, Sigma) followed by the above immunostaining procedures.

To quantify blood vessel density, we used projected confocal images at 40x magnification (Nikon C1) followed by NIH ImageJ automated processing. At least three sections from each mouse were used for ImageJ quantification. Ten-micron-thick optical sections from confocal *z*-stack images were projected into a flattened image. The parameter setting of z-stack confocal imaging is: total optical thickness, 10μm, step size, 0.5 μm, total number optical slices, 21. The volume-rendered confocal images were subsequently imported to NIH ImageJ 1.46r for quantifying Laminin-positive blood vessel density using a customer-defined Macro program. The total area and Laminin-occupying area were derived and from ImageJ and exported to Microsoft Excel for calculating the percent of Laminin-occupying area among assessed total CNS area.

### Complementary RNA (cRNA) probe preparation and dual fluorescence mRNA In Situ hybridization (ISH)

We employed the PCR and in vitro transcription to prepare cRNA probes targeting Vegfa and Plp^58^. Targeted sequences of Vegfa and Plp were generated by PCR. The primers used were: *Vegfa*-Forward: GGATATGTTTGACTGCTGTGGA; *Vegfa*-Reverse: AGGGAAGATGAGGAAGGGTAAG; *Plp*-Forward: GGGGATGCCTGAGAAGGT; *Plp*-Reverse: TGTGATGCTTTCTGCCCA. We added the T7 (GCGTAATACGACTCACTATAGGG) and SP6 (GCGATTTAGGTGACACTATAG) promoter sequences to the 5’ of the forward and reverse primers, respective. The SP6 and T7 promoter sequences are recognized by the SP6 and T7 RNA polymerase, respectively, in the subsequence in vitro transcription. PCR products of Vegfa and Plp amplification were used as DNA templates to transcribe into Vegfa and Plp cRNA probes in vitro using SP6 RNA polymerase. T7 RNA polymerase-mediated transcription of RNA was used as negative control. DIG-UTP or FITC-UTP was used to generate DIG- or FITC-labeled cRNA probes.

Single or dual mRNA ISH was done using our previous protocols^58^. Frozen sections of 14 μm thick were used. The concentration of cRNA probe we used was 100ng/100μl hybridization buffer. Hybridization was conducted at 65°C for 18–20 h. After hybridization, sections were treated with 10 μg/ml RNase A to eliminate nonspecific cRNA binding. For single mRNA ISH using DIG-labeled cRNA probes, DIG was recognized by alkaline phosphatase (AP)-conjugated anti-DIG (#11093274910, Sigma) antibody and DIG signals were visualized by the NBT/BCIP (#72091, Sigma) method. For dual fluorescence mRNA ISH (Vegfa and Plp), FITC-labeled Plp cRNA probe and DIG-labeled Vegfa cRNA probe were applid to frozen sections simultaneously during the hybridization step. The FITC signals were visualized by tyramide signal amplification (TSA) fluorescence system (#NEL747A,Perkin Elmer) according to the manufacturer’s instructions using horseradish peroxidase (HRP)-IgG Fraction Monoclonal Mouse Anti-Fluorescein (#200-032-037,Jackson ImmunoResearch). DIG signals were visualized by a HNPP fluorescent kit (#11758888001, Sigma) according to the manufacturer’s instructions using AP-conjugated anti-DIG Fab^2^ antibody (#11093274910, Sigma).

### Western Blot

Protein concentration was assessed by BCA protein assay kit (#23225, Thermo Fisher Scientific). Twenty microgram protein lysates were separated on AnykD Mini-PROTEAN TGX precast gels (#4569035, BIO-RAD) or 10% Mini-PROTEAN TGX precast gels (#4561035, BIO-RAD). The proteins were transferred onto 0.2μm nitrocellulose membrane (#1704158, BIO-RAD) by Trans-blot Turbo Transfer system (#1704150, BIO-RAD). The membranes were blocked with 5% BSA (#9998, Cell signaling) for 1h at room temperature and were incubated overnight with primary antibodies (Supplementary Table 2) at 4 °C. The membranes were washed 3 times with 10 mm Tris-HCl (pH7.5) containing 150 mM NaCl and 0.1% Tween-20 (TBST) and were incubated with horseradish peroxidase-conjugated goat anti-rabbit (31460, RRID: AB_228341, Thermo Fisher Scientific) or anti-mouse (31430, RRID: AB_228307, Thermo Fisher Scientific) for 1h at room temperature. After incubation, the membranes were washed 3 times with TBST. Specific binding was detected using Western Lightening Plus ECL (NEL103001EA, Perkin Elmer). NIH Image J was used to quantify protein expression levels by analyzing the scanned grey-scale films

### RNA preparation, reverse transcription and real-time quantitative PCR (RT-qPCR)

Total RNA was extracted by using Qiagen RNeasy for lipid tissues (74804, Qiagen) with additional on-column DNase I digestion to remove genomic DNA contamination. The quality and quantity of RNAs were analyzed by the Nanodrop one^C^ microvolume UV-Vis Spectrophotometer (ND-ONEC-W, Thermo Fisher Scientific). cDNA was synthesized by Qiagen Omniscript RT Kit (205111, Qiagen). The relative mRNA level of indicated genes was normalized to that of the internal control Hsp90 and calculated by the equation 2^^(Ct(cycle threshold) of Hsp90 - Ct of indicated genes)^. The gene expression levels in control groups were normalized to 1. RT-qPCR was conducted by QuantiTect SYBR® Green PCR Kit(204145, QIAGEN) approaches on Agilent MP3005P thermocycler. The qPCR primers used in the study were listed in Supplementary Table 3.

### Statistical Analyses

Quantification was performed by blinded observers. All measurements were taken from distinct mice and quantitative data are presented as means ± standard deviation (s.d.). We used scatter dot plots to present the quantification data throughout our manuscript. Each dot (circle, square, or triangle) in the scatter dot plots represents one mouse or one independent experiment. Shapiro-Wilk approach was used for testing data normality. F test was used to compare the equality of variances of two groups whereas Browne-Forsythe test was used for comparing the equality of variances of three or more groups. The statistical methods were described in the figure legends and P value was presented in each graph. For unpaired, two-tailed Student’s *t* test, t value and degree of freedom (df) were presented as t_(df)_ in figure legends. Welch’s correction was used for Student’s *t* test if the variances of two groups were unequal after Browne-Forsythe test. For comparisons among three or more groups with equal variances (tested by Browne-Forsythe approach), ordinary one-way ANOVA was used followed by Tukey’s multiple comparisons, otherwise Welch’s ANOVA was used followed by unpaired *t* test with Welch’s correction. In ordinary ANOVA, the F ratio and DFn and DFd was presented as F_(DFn, DFd)_ in the figure legends where DFn stands for degree of freedom numerator and DFd for degree of freedom of denominator. In Welch’s ANOVA, the Welch’s F ratio, W and DFn and DFd was presented as W_(DFn, DFd)_ in the figure legends. All data plotting and statistical analyses were performed using GraphPad Prism version 8.0. P value less than 0.05 was considered as significant, whereas greater than 0.5 was assigned as not significant (ns).

## Acknowledgements

This study was supported by the grants funded by NIH/NINDS (R21NS109790, R01NS094559, and R21NS093559 to F.G.) and Shriners Hospitals for Children (86100, 85200-NCA16, 85107-NCA-19 to F.G., 84307-NCAL to S.Z.).

## Contributions

F.G. conceived the original idea, led the project, and wrote the manuscript. S.Z, B.K. conceived the idea, designed and performed experiments, analyzed data, and wrote the manuscript. X.Z. X.G. Y.W. Z.L. P.P. K.F, A.W. analyzed data and edit the manuscript. All the authors provided feedback and comments on the manuscript.

