## Supplementary information for "Glial type specific regulation of CNS angiogenesis by HIFα-activated different signaling pathways"

Zhang, Kim et al.,

14 supplementary figures and 3 supplementary tables.

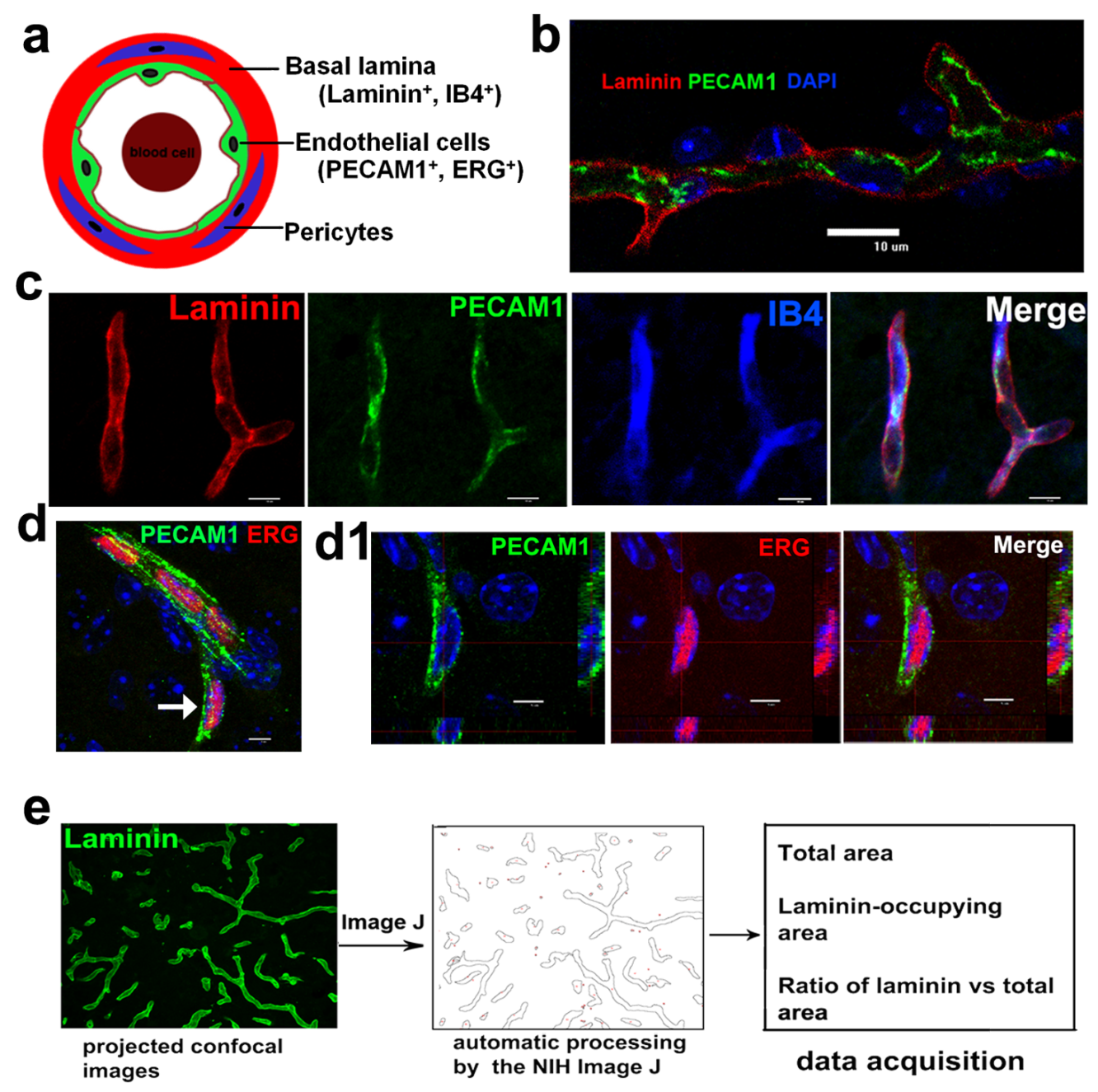

**Supplementary Figure 1 - Semi-automated quantification of blood vessel density.**

**a,** simplified diagram depicting the expression of the endothelial cell markers platelet/endothelial cell adhesion molecule 1 (PECAM1) and ETS-related gene (ERG) and of blood vessel basement membrane markers Laminin and isolectin-B4 (IB4). **b**, single optic slice of confocal images showing Laminin and PECAM1 expression in the blood vessels. DAPI, nuclei counterstaining. **c**, projected confocal images showing Laminin, PECAM1 and IB4 expression in the blood vessels. **d-d1**, projected confocal images (**d**) showing PECAM1 and ERG expression in the blood vessels. Blue is the DAPI nuclei counterstaining. The endothelial cell pointed by the arrow is shown in **d1** at higher magnification and orthogonal views. Scale bars, 10 μm, applied to all. **e,** semi-automated approach for quantifying blood vessels. Projected confocal images of 10 μm optical section are converted to binary images by the NIH Image J using the programed Macro functions. Laminin^+^ blood vessel area and total area are retrieved to calculate the ratio of blood vessel-occupying area to total area measured.

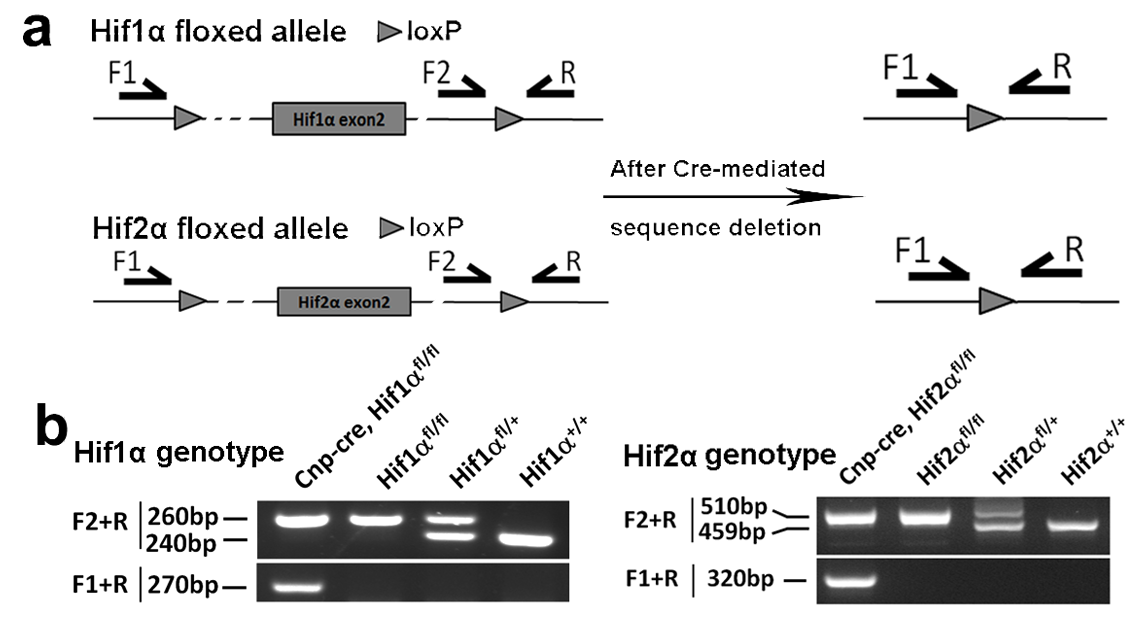

**Supplementary Figure 2 – PCR primer design and detection of Hif1α and Hif2α gene deletion.**

**a,** primer pair of F2/R is used for genotyping Hif1α and Hif2α genes, respectively and primer pair of F1/R is used for detecting the sequence deletion of Hif1α and Hif2α genes. **b**, After Cre-mediated gene deletion, PCR amplification by primer pair F1/R generates a 270bp product in the genome of *Cnp-Cre: Hif*1α^fl/fl^ and a 320bp product of *Cnp-Cre: Hif*2α^fl/fl^ transgenic mice.

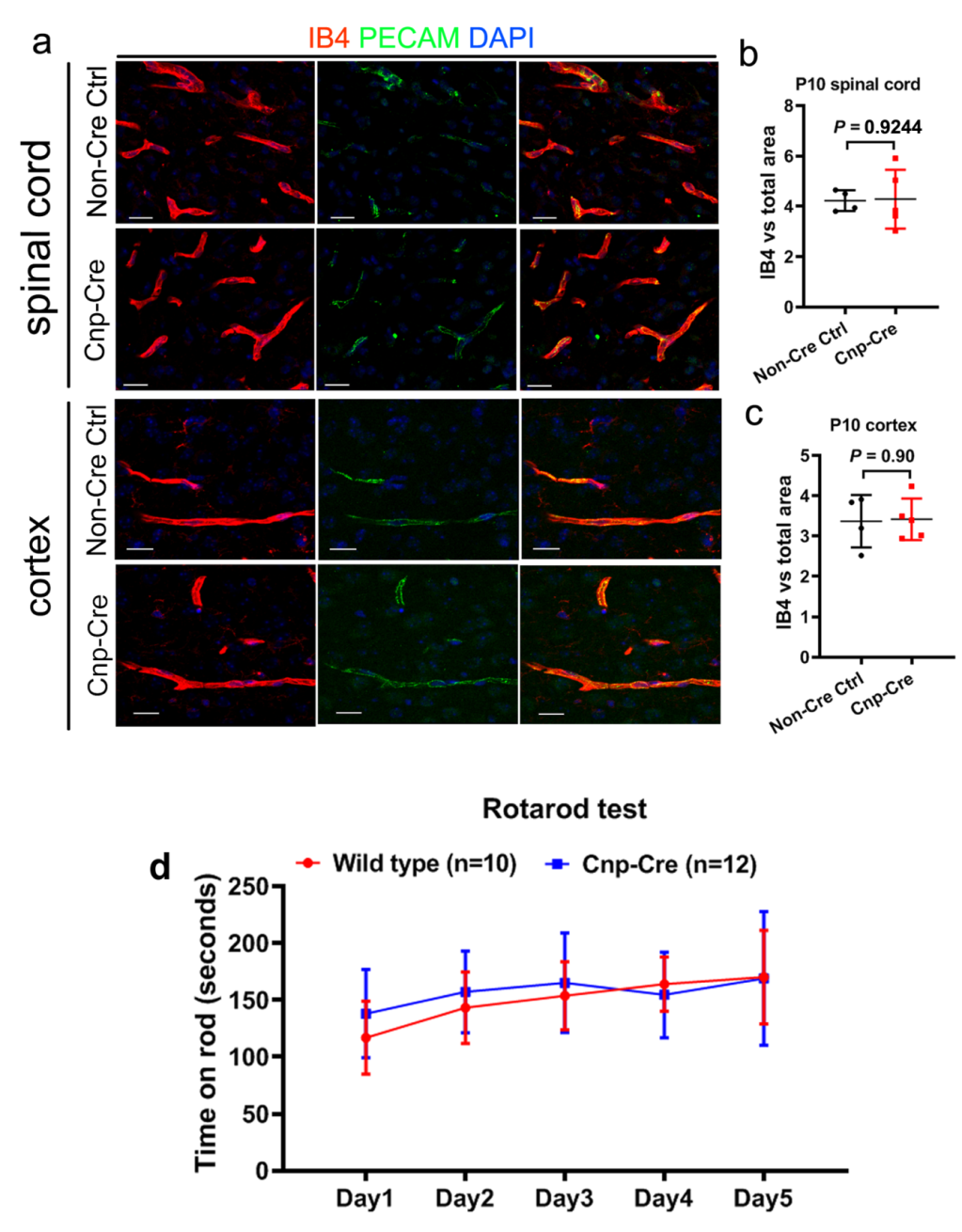

**Supplementary Figure 3 - Cnp-Cre has no effect on CNS angiogenesis and animal behaviors.**

**a,** representative confocal images of IB4, PECAM1, and DAPI immunostaining in the spinal cord and cerebral cortex of Non-Cre Ctrl (wild type) and Cnp-Cre transgenic mice. Scale bars = 20 µm. **b**, percent of IB4-positive area among total area assessed in the spinal cord at P10. Two-tailed Student’s *t* test, t_(7)_=0.0983. **c**, percent of IB4-positive area among total area assessed in the cerebral cortex at P10. Two-tailed Student’s *t* test, t_(7)_=0.1239. **d**, time retention on rod during a 5-day course of accelerating Rotarod test starting at one month old. Two-way ANOVA, *F*_(1, 20)_ = 0.2741, *P* = 0.6081 for genotype. Source data of **b-d** are provided as a Source Data file.

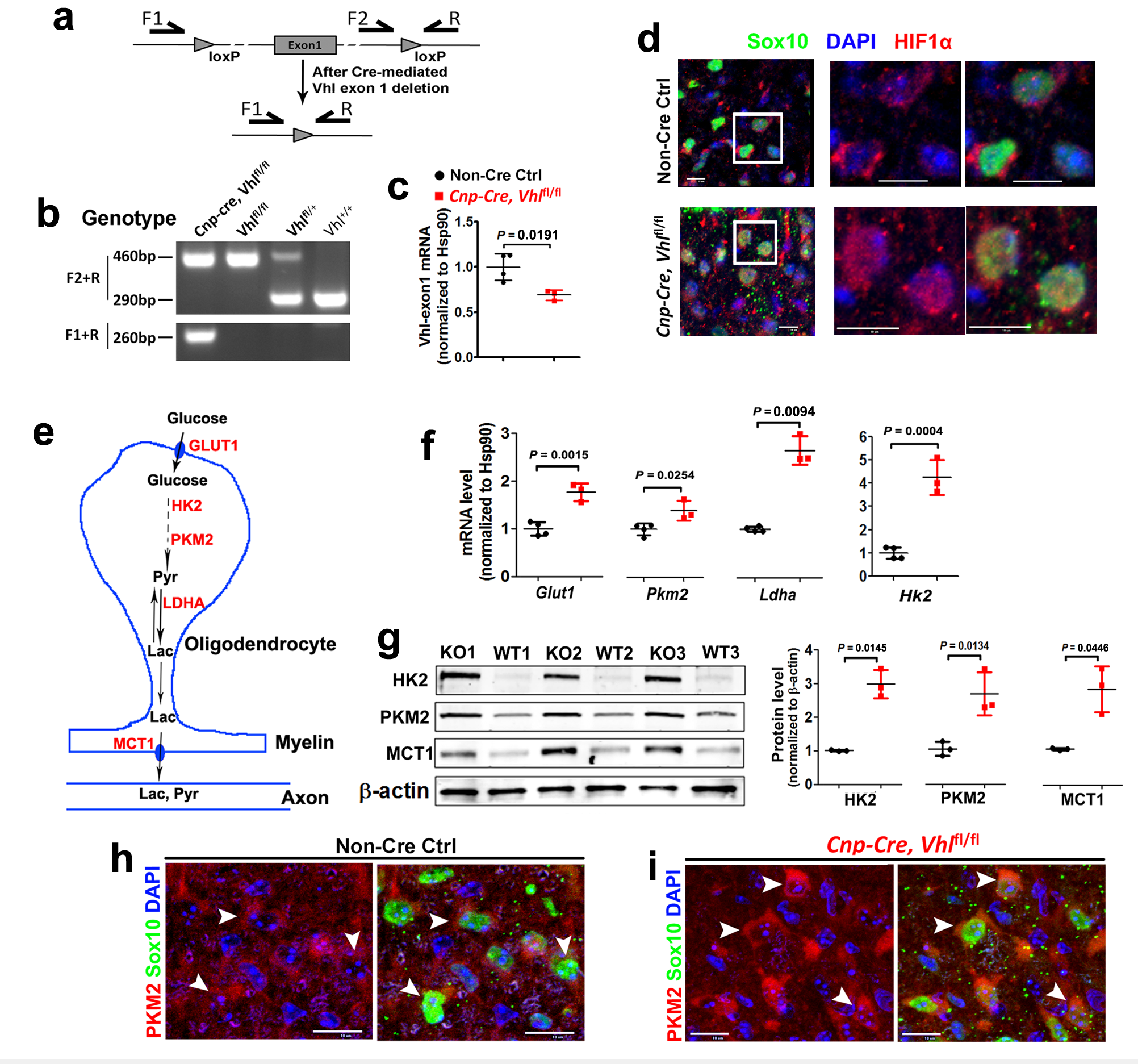

**Supplementary Figure 4 – Oligodendroglial VHL disruption stabilizes HIFα signaling pathway.**

**a-b**, primer design (**a**) and PCR detection (**b**) of *Vhl* gene deletion. Primer pair of F2/R is for genotyping *Vhl* gene. After Cre-mediated deletion, PCR amplification by primer pair F1/R generates a 260bp product in the genome of *Cnp-Cre: Vhl*^fl/fl^ transgenic mice. **c**, RT-qPCR quantification of the mRNA level transcribed from *Vhl* exon1 in the spinal cord at P8. Two-tailed Student’s t test, t_(5)_ = 3.408. **d**, HIF1α and Sox10 immunohistochemistry showing HIF1α protein is diffusely located in cytoplasm in non-Cre Ctrl (left panels, arrowheads) but concentrated in the nuclei of Sox10^+^ oligodendroglial lineage cells of *Cnp-Cre:Vhl*^fl/fl^ spinal cord (right panels, arrowheads) at P8. Scale bar: 10 μm. **e**, diagram depicting glucose metabolic pathway through glycolysis. GLUT1 glucose transporter 1, HK2 hexokinase 2, PKM2 pyruvate kinase isoform M, LDHA lactate dehydrogenase A, MCT1 monocarboxylate transporter 1. Pyr pyruvate, Lac lactate. Glycolytic pathway is one of the established pathways activated by HIFα, and the key enzymes in glycolytic pathway are direct downstream target of HIFα. **f**, RT-qPCR quantification of mRNA levels. mRNA levels are normalized to the internal control HSP90 and designated as 1 in non-Cre control spinal cord. Two-tailed Student’s *t* test, t_(5)_ = 6.264 *Glut1*, t_(5)_ = 3.148 *Pkm2*, t_(5)_ = 8.282 *Hk2*. Two-tailed Student’s *t* test with Welch’s correction, t_(2.1)_ = 9.436 *Ldha.* **g**, Western blot images and quantifications. Protein levels are normalized to the internal loading control β-action and designated as 1 in non-Cre control spinal cord. Two-tailed Student’s t test, Welch’s corrected t_(2.01)_ = 8.150 HK2, t_(4)_ = 4.231 PKM2, Welch’s corrected t_(2.011)_ = 4.550 MCT1. **h-i**: double immunohistochemistry showing that PMK2 is expressed in Sox10^+^ oligodendroglial lineage cells in non-Cre control spinal cord (**h,** arrowheads) and is upregulated in *Cnp-Cre:Vhl*^fl/fl^ mice (**i**, arrowheads). All data are from spinal cord at P8 except for Panel **g** at P14. Black circles represent data from non-Cre control and red squares from *Cnp-Cre:Vhl*^fl/fl^. Scale bar: 10 μm. Source data of **b-c, f-g** are provided as a Source Data file.

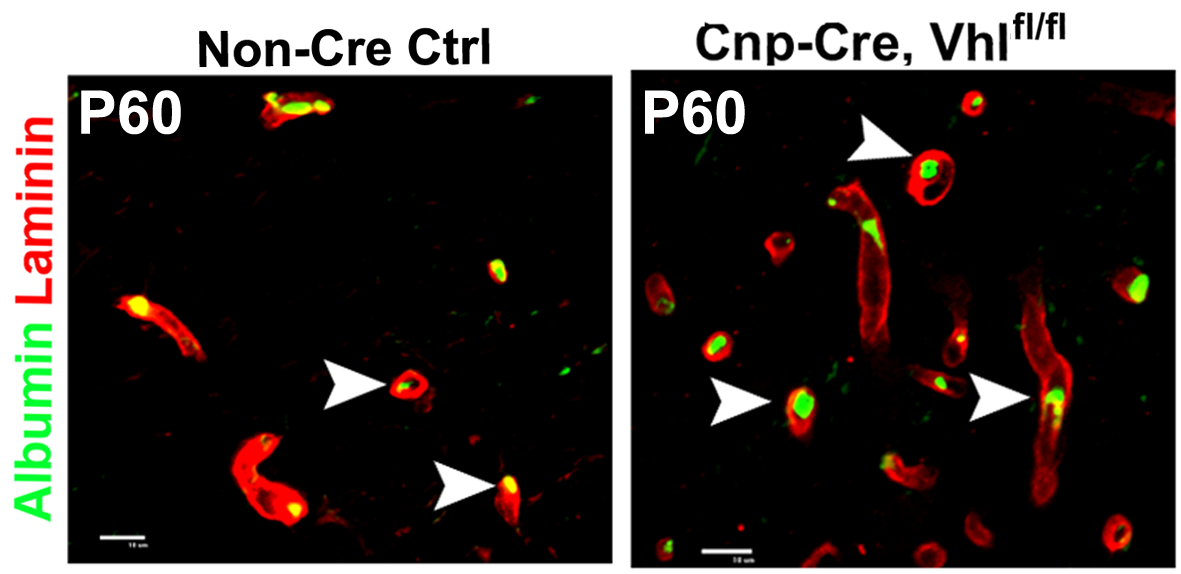

**Supplementary Figure 5 – Stabilizing oligodendroglial HIFα does not affect the integrity of the blood brain (spinal cord) barrier.**

The blood-borne macromolecules such as albumin and immunoglobin (Ig) cannot cross over the blood vessels and spread into the CNS parenchymal tissues due to the function of the blood brain barrier. Immunohistochemistry showed that albumin was restricted to the lumen of the blood vessels (arrowheads) in the spinal cord of *Cnp-Cre, Vhl*^fl/fl^ and non-Cre control mice at P60 though the blood vessel density was elevated. Scale bars = 10 μm.

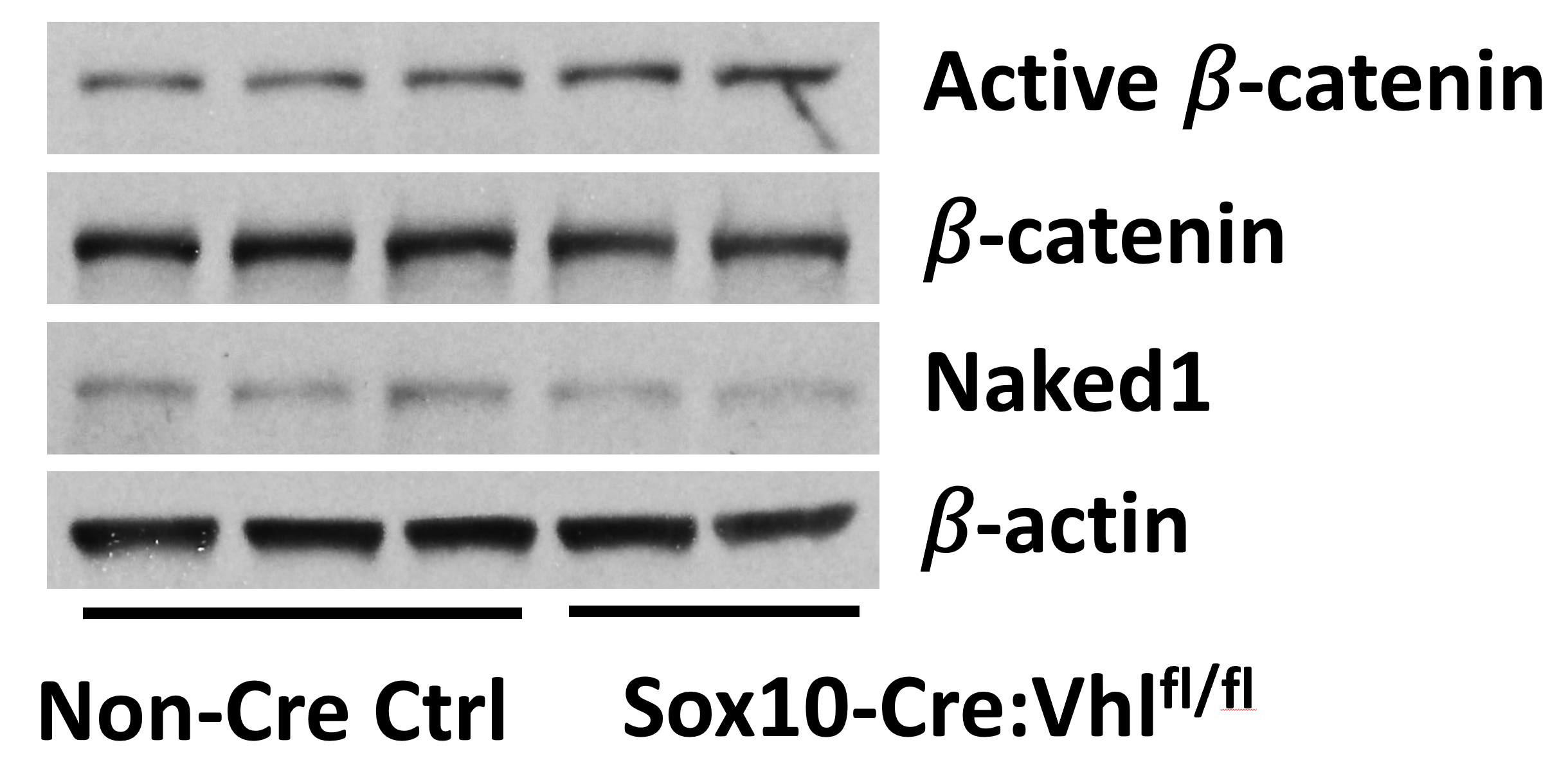

**Supplementary Figure 6 – Wnt/β-catenin signaling activity is not changed in the spinal cord of Sox10-Cre:Vhl^fl/fl^ mutants.**

Western blot images showing Wnt signaling active β-catenin, total β-catenin, Wnt target gene Naked1, and internal loading control β-actin were similar between Sox10-Cre:Vhl^fl/fl^ mutants (n=2) and Non-Cre control mice (n=3) in the spinal cord at P5. One spinal cord sample from Sox10-Cre:Vhl^fl/fl^ group was lost during protein sample preparation. Source data are provided as a Source Data file.

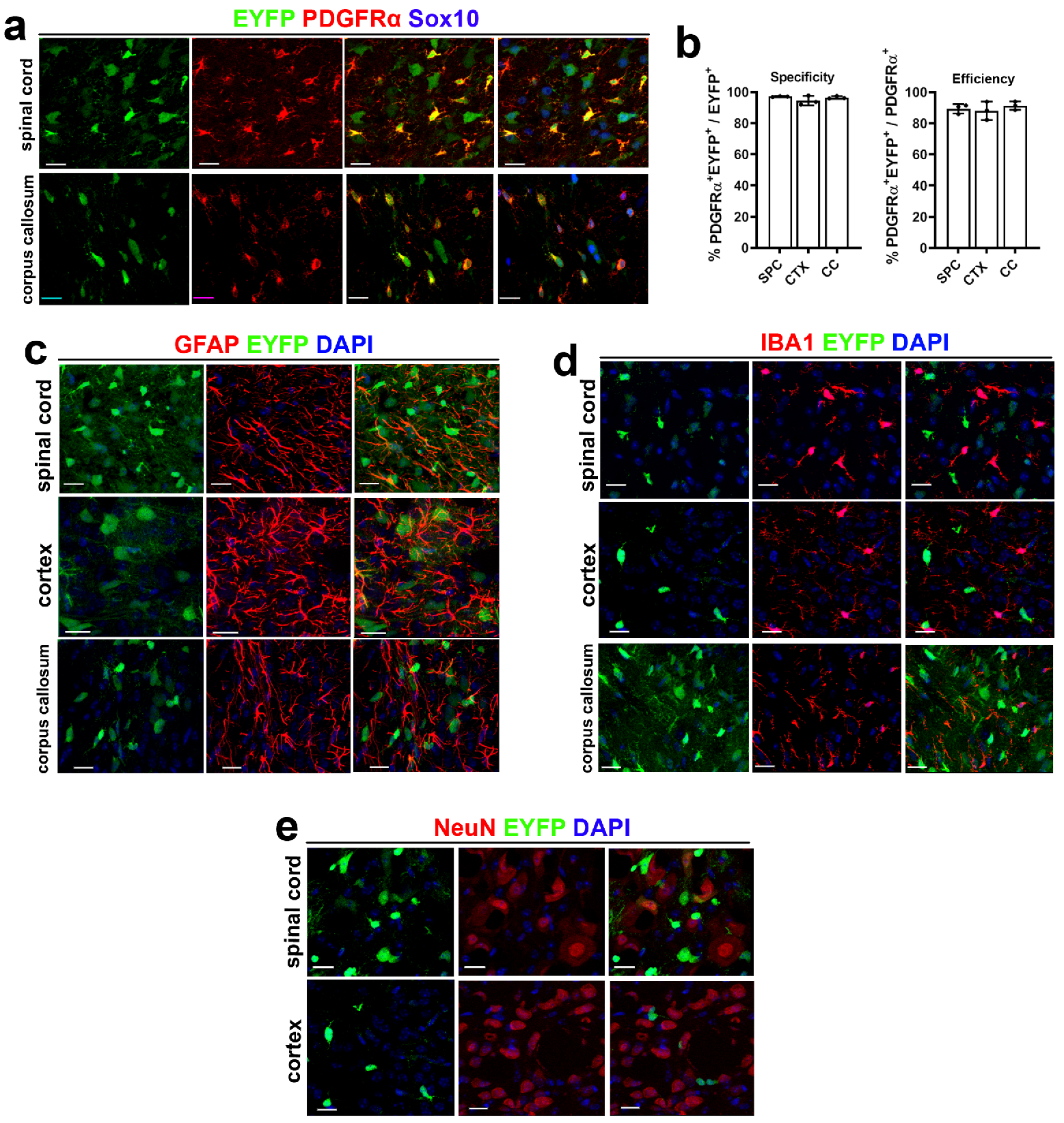

**Supplementary Figure 7 - Recombination efficiency and specificity of Pdgfrα-CreER^T2^ in the CNS.**

Tamoxifen was administered to Pdgfrα-CreER^T2^:Rosa26-EYFP double transgenic reporter mice at P6 and P7 and the brain and spinal cord were harvested for analysis at P14. **a**, immunohistochemistry of EYFP, PDGFRα (OPC marker), and Sox10 (pan-oligodendroglial lineage marker) in the spinal cord and corpus callosum. Scale bars = 20 µm. **b**, Pdgfrα-CreER^T2^-mediated recombination efficiency (percent of PDGFRα^+^EYFP^+^ cells among PDGFRα^+^ cells) and OPC specificity (percent of PDGFRα^+^EYFP^+^ cells among EYFP^+^ cells). SPC, spinal cord; CTX, cerebral cortex; CC, corpus callosum. **c-e**, immunostaining of EYFP, DAPI, and astrocytic marker GFAP (**c**), microglial marker IBA1 (**d**), and neuronal marker NeuN (**e**) in different areas of the CNS. Scale bars = 20 µm. Source data of **b** are provided as a Source Data file.

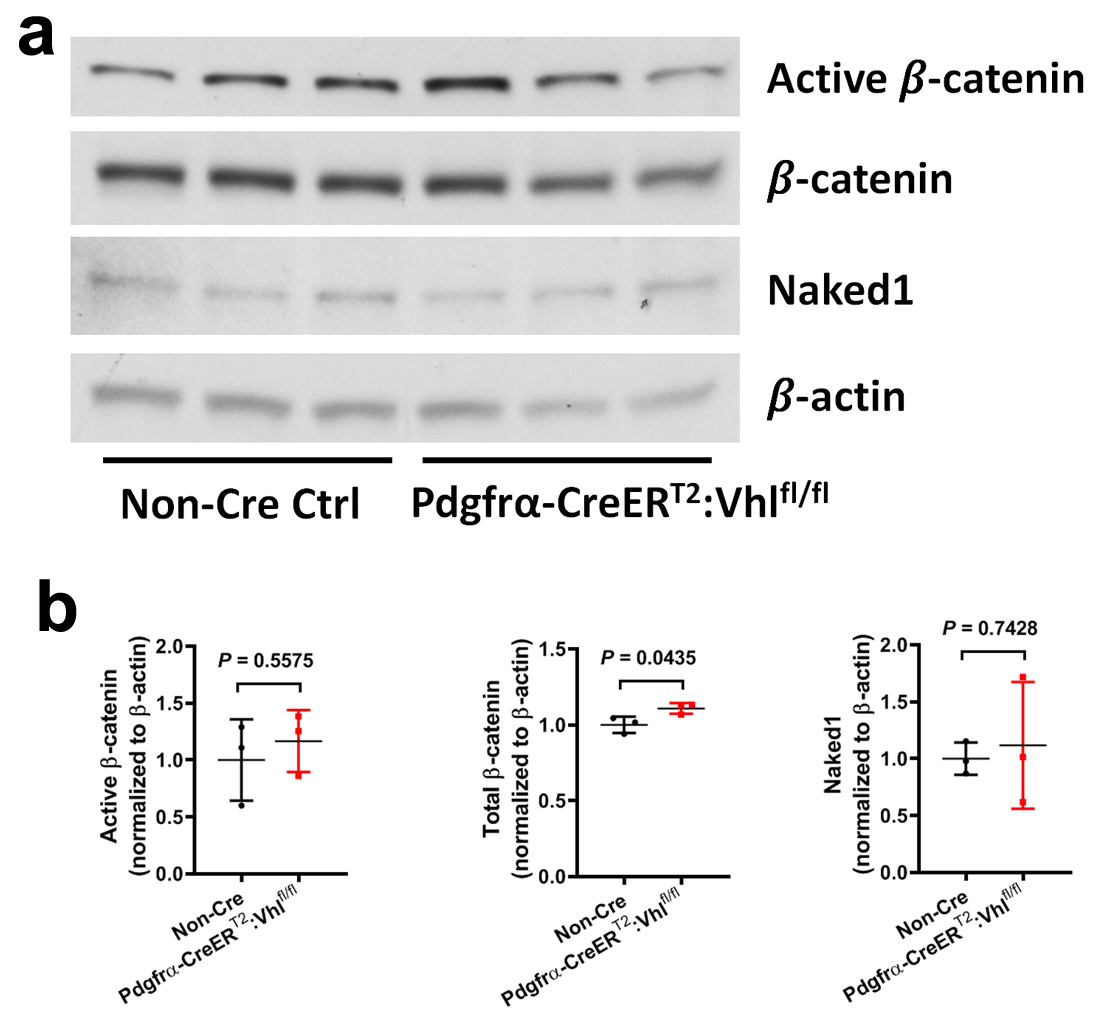

**Supplementary Figure 8 – Wnt/β-catenin signaling activity is not changed in the spinal cord of Pdgfrα-CreER^T2^:Vhl^fl/fl^ mutants.**

**a-b**, Western blot images showing Wnt signaling active β-catenin, total β-catenin, Wnt target gene Naked1, and internal loading control β-actin. **b**, quantification of blotting signals. Two-tailed Student’s t test, t(4) = 0.6392 active β-catenin; t(4) = 2.914 total β-catenin; t(4) = 0.3517 Naked1. Tamoxifen was administered to Pdgfrα-CreER^T2^:Vhl^fl/fl^ and non-Cre control animals at P6 and P7 and tissue was harvested at P14. Source data are provided as a Source Data file.

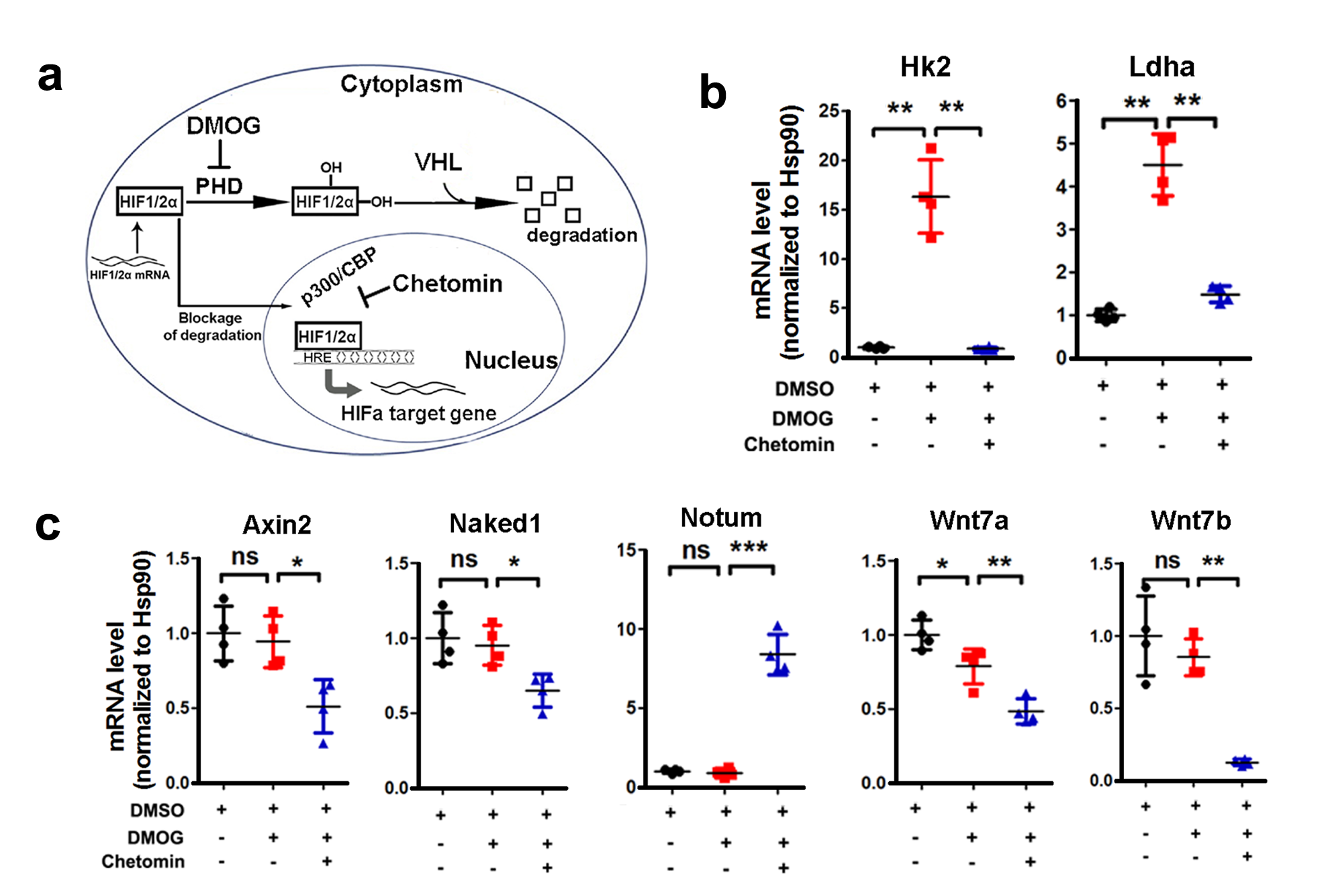

**Supplementary Figure 9 - Pharmacologically stabilizing HIFa does not activate Wnt signling in primary OPCs.**

**a**, schematic diagram depicting HIFα signaling and its modulation by small compounds. Dimethyloxalylglycine (DMOG) stabilizes HIFα signaling by inhibiting prolyl-hydroxylase (PHD) whereas Chetomin inhibits HIFα signaling by disrupting the function of the common co-activator P300/CBP, thus preventing its binding of P300/CBP to HIFα. Purified cortical OPCs from neonatal mouse forebrains are pre-incubated with Chetomin (100nM) or DMSO control for 2 hrs, then treated with DMOG (1mM) in the presence of Chetomin (100nM) or DMSO control for 7 hrs and mRNA of indicated genes is quantified by RT-qPCR. N=4 independent experiments. **b**, RT-qPCR assay of the mRNA levels of the HIFα target genes Hk2 and Ldha. Welch’s ANOVA followed by unpaired *t* test with Welch’s correction, ** *P* < 0.01. *W*_(2, 5.310)_ = 30.52, *P* = 0.0012 *Hk2*, *W*_(2, 5.263)_ = 43.89, *P* = 0.0005 *Ldha*. **c**, RT-qPCR assay of the mRNA levels of Wnt/β-catenin target genes *Axin2*, *Naked1*, *Notum*, and the ligands *Wnt7a* and *Wnt7b*. One-way ANOVA followed by Tukey’s multiple comparisons, * *P* < 0.05, ** *P* < 0.01, ns, not significant. *F*_(2, 9)_ = 9.064, *P* = 0.007 *Axin2*; *F*_(2, 9)_ = 7.358, *P* = 0.0128 *Naked1*; *F*_(2, 9)_ = 25.62, *P* = 0.0002 *Wnt7a*. Welch’s ANOVA followed by unpaired *t* test with Welch’s correction, ** *P* < 0.01, *** *P* < 0.001, ns, not significant. *W*_(2, 4.696)_ = 58.98, *P* = 0.0005 *Notum*; *W*_(2, 4.138)_ = 69.71, *P* = 0.0007 *Wnt7b*. Note that the activity of Wnt signaling was downregulated in the presence of Chetomin presumably due to the disruption of the interaction of P300/CBP with the Wnt effector TCF/LEF1 complex and/or to possible off-target effects. Source data of **b-c** are provided as a Source Data file.

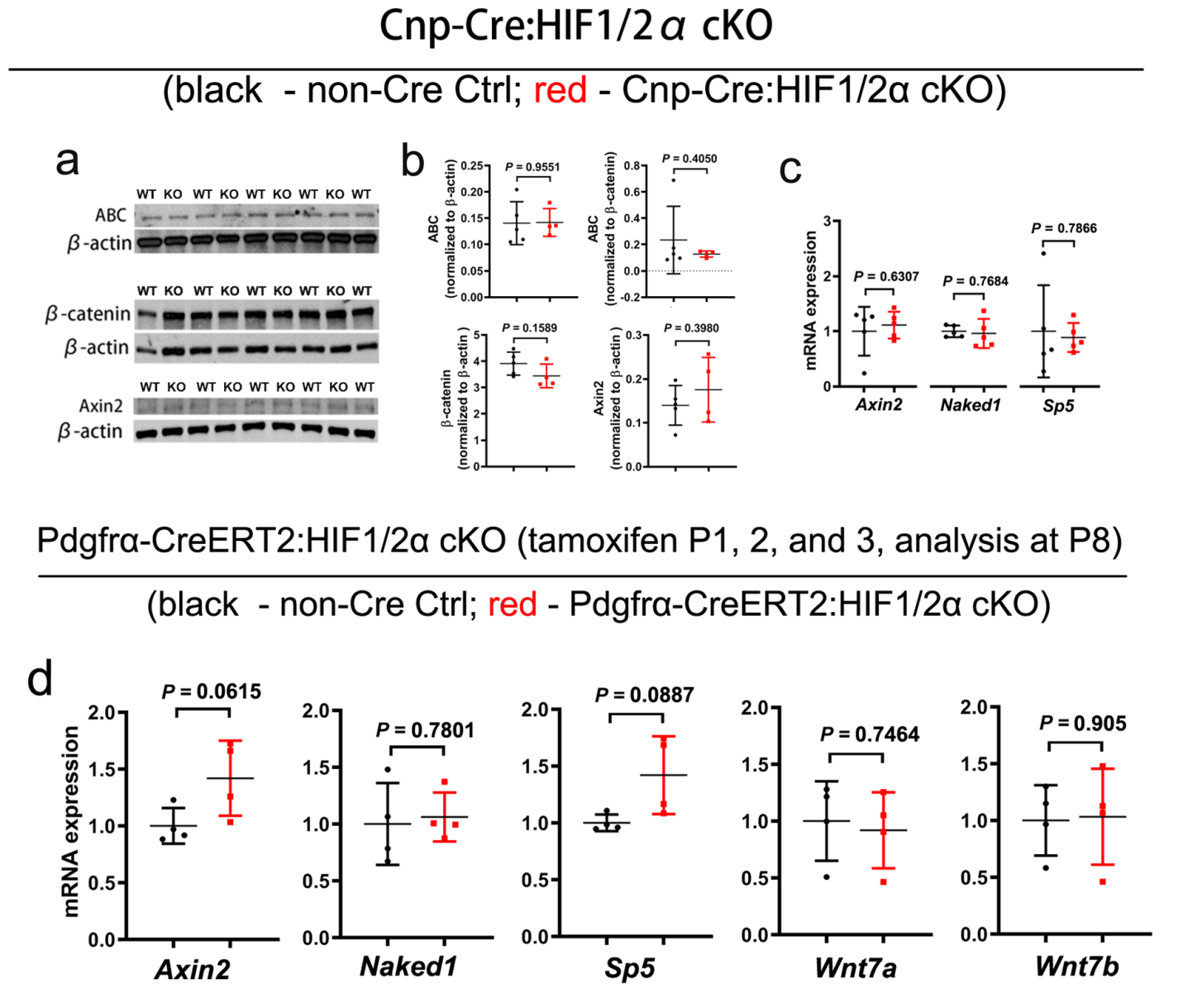

**Supplementary Figure 10 – Oligodendroglial HIFα deletion does not affect Wnt/β-catenin signaling activity in the CNS.**

**a-b,** Western blot and quantification of active β-catenin (ABC), total β-catenin, and Axin2 in the spinal cord of P14 Cnp-Cre, Hif1α^fl/fl^, Hif2α^fl/fl^ mutants (Cnp-Cre:HIF1/2α cKO) and non-Cre littermate controls. Two-tailed Student’s *t* test. t_(7)_ = 0.0584 ABC normalized to loading control β-actin; Welch’s-corrected t_(4.081)_ = 0.9279 ABC normalized to total β-catenin; t_(7)_ = 1.577 β-catenin normalized to β-actin; t_(7)_ = 0.900 Axin2 normalized to β-actin. **c**, RT-qPCR assay of the mRNA levels of Wnt/β-catenin signaling target genes in the spinal cord of P14 Cnp-Cre:HIF1/2a cKO and non-Cre Ctrl. Two-tailed Student’s *t* test. t_(8)_ = 0.4998 *Axin2*; t_(8)_ = 0..3047 *Naked1*; Welch’s-corrected t_(4.7888)_ = 0.2863 *Sp5*. **d**, RT-qPCR assay of the mRNA levels of Wnt/β-catenin signaling target genes and Wnt7a/b in the spinal cord of P8 Pdgfrα-CreER^T2^, Hif1α^fl/fl^, Hif2α^fl/fl^ mutants (Pdgfrα-CreER^T2^:HIF1/2a cKO) and non-Cre Ctrl that had been treated with tamoxifen at P1, P2, and P3. Two-tailed Student’s *t* test. t_(6)_ = 2.925 *Axin2*; t_(6)_ = 0.292 *Naked1*; Welch’s-corrected t_(3.274)_ = 2.399 *Sp5*; t_(6)_ = 0.3383 *Wnt7a*; t_(6)_ = 0.1244 *Wnt7b.* Source data are provided as a Source Data file.

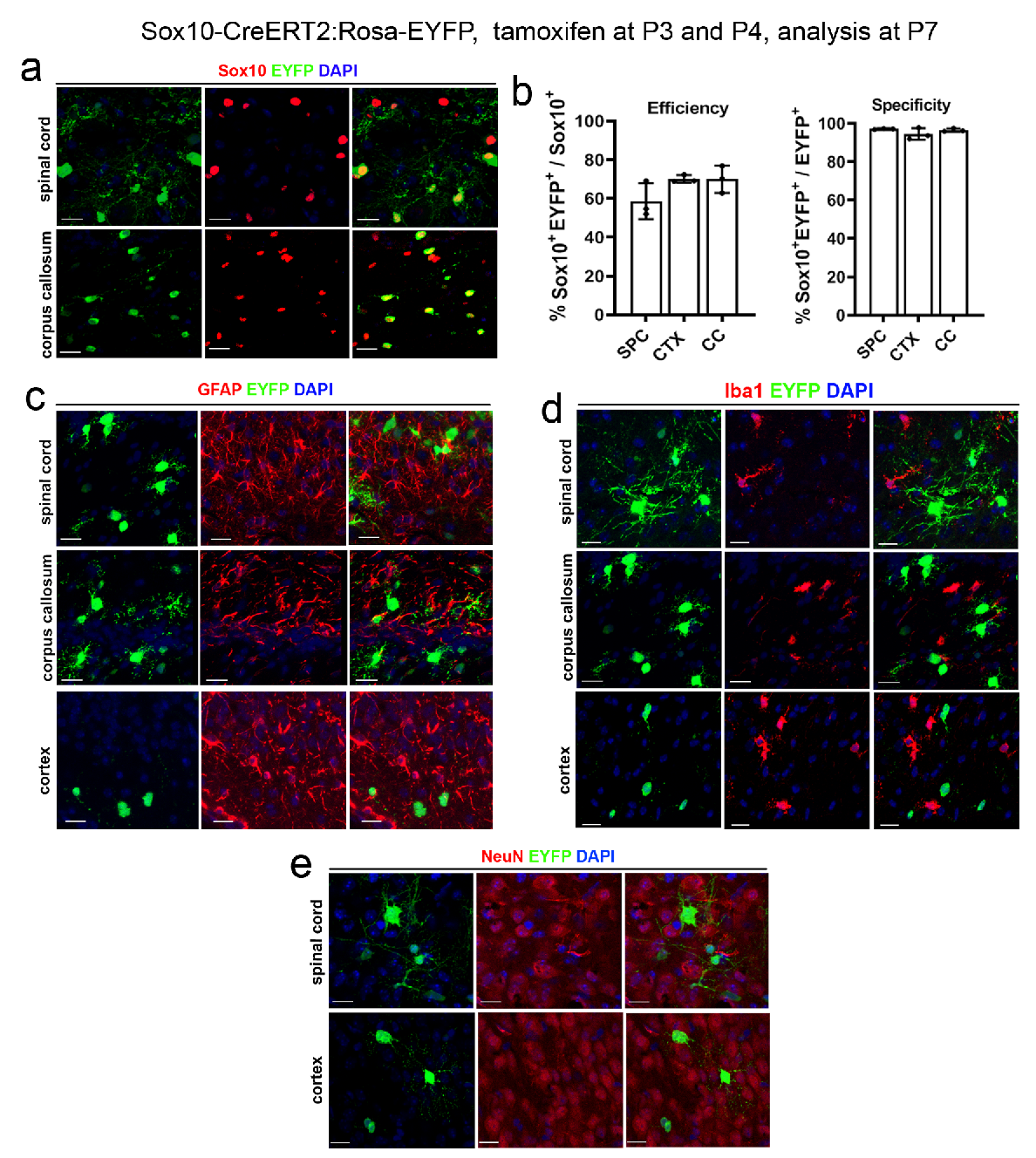

**Supplementary Figure 11 - Recombination efficiency and specificity of Sox10-CreER^T2^ in the CNS.**

**a**, immunohistochemistry of EYFP, Sox10 (pan-oligodendroglial lineage marker), and nuclear DAPI in the spinal cord and corpus callosum. Scale bars = 20 µm. **b**, Sox10-CreER^T2^-mediated recombination efficiency (percent of Sox10^+^EYFP^+^ cells among Sox10^+^ cells) and oligodendroglial lineage specificity (percent of Sox10^+^EYFP^+^ cells among EYFP^+^ cells). SPC, spinal cord; CTX, cerebral cortex; CC, corpus callosum. **c-e**, immunostaining of EYFP, DAPI, and astrocytic marker GFAP (**c**), microglial marker IBA1 (**d**), and neuronal marker NeuN (**e**) in different areas of the CNS. Scale bars = 20 µm. Tamoxifen was administered to Sox10-CreER^T2^:Rosa26-EYFP double transgenic reporter mice at P3 and P4 and the brain and spinal cord were harvested for analysis at P7. Source data of **b** are provided as a Source Data file.

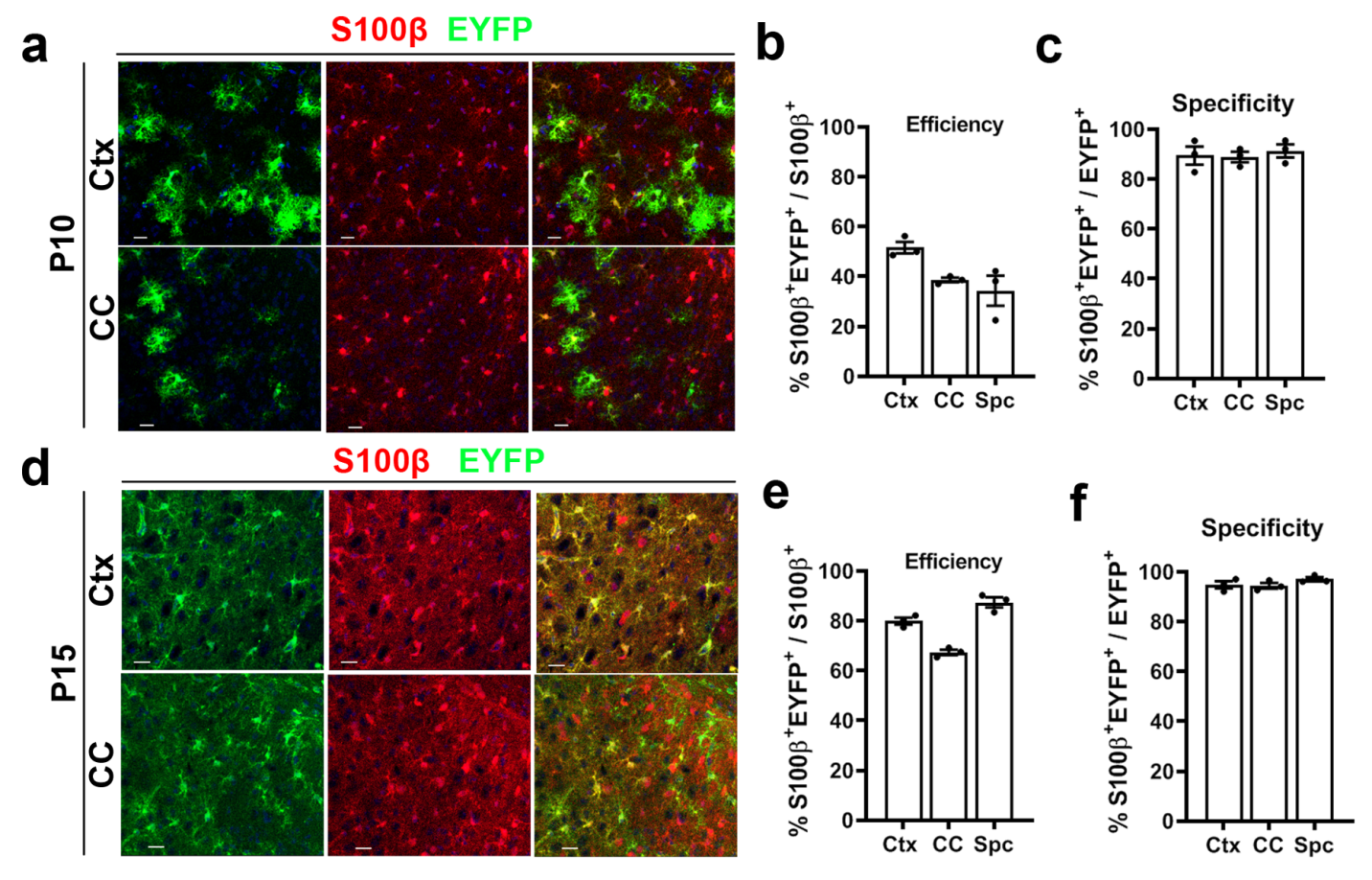

**Supplementary Figure 12 - Recombination efficiency and specificity of mGfap-Cre in the early postnatal CNS.**

mGfap-Cre mice were crossed with reporter mice Rosa26-EYFP to generate mGfap-Cre:Rosa26-EYFP hybrid mice and the brain and spinal cord were harvested for analysis at P10 and P15. **a**, representative images of EYFP and astrocytic marker S100β in the cerebral cortex (Ctx) and corpus callosum (CC) at P10. Scale bars = 20 µm**. b-c**, mGfap-Cre-mediated recombination efficiency (percent of S100β^+^EYFP^+^ cells among S100β^+^ cells) and astrocyte specificity (percent of S100β^+^EYFP^+^ cells among EYFP^+^ cells). Spc, spinal cord. **d**, representative images of EYFP and astrocytic marker S100β in the cerebral cortex (Ctx) and corpus callosum (CC) at P15. Scale bars = 20 µm. **e-f**, Cre-mediated recombination efficiency (percent of S100β^+^EYFP^+^ cells among S100β^+^ cells) and astrocyte specificity (percent of S100β^+^EYFP^+^ cells among EYFP^+^ cells). Spc, spinal cord. Source data of **b-c, e-f** are provided as a Source Data file.

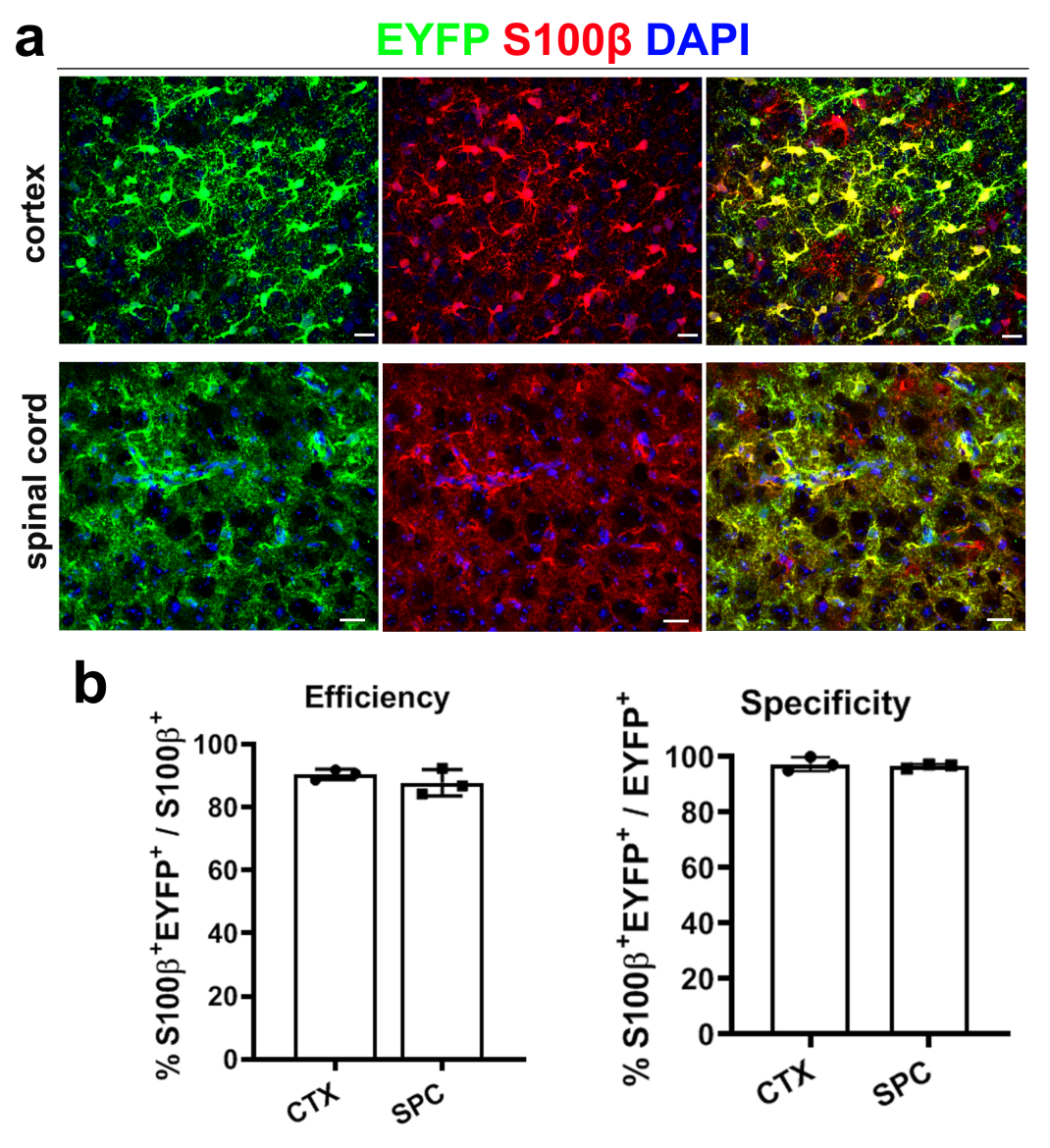

**Supplementary Figure 13 - Recombination efficiency and specificity of Aldh1l1-CreERT2 in the early postnatal CNS.**

Tamoxifen was administered to Aldh1l1-CreER^T2^:Rosa26-EYFP double transgenic reporter mice at P1, P2, and P3 and the brain and spinal cord were harvested for analysis at P8. **a**, immunohistochemistry of EYFP, S100β (astrocyte marker), and DAPI in the cerebral cortex and spinal cord. Scale bars = 20 µm. b, Aldh1l1-CreER^T2^-mediated recombination efficiency (percent of S100β^+^EYFP^+^ cells among S100β^+^ cells) and astrocyte specificity (percent of S100β^+^EYFP^+^ cells among EYFP^+^ cells). SPC, spinal cord; CTX, cerebral cortex. Source data of **b** are provided as a Source Data file.

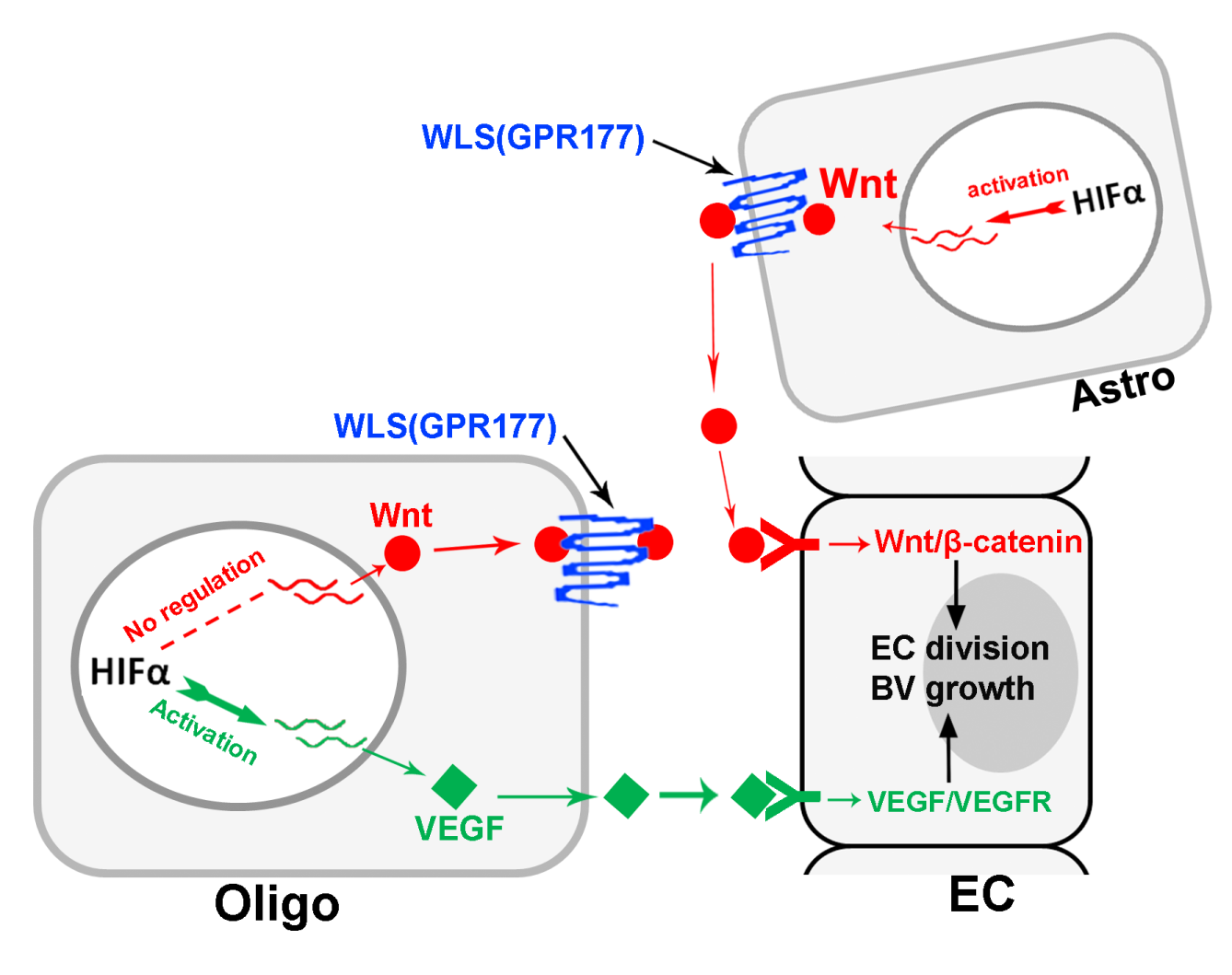

**Supplementary Figure 14 – An alternative working model of glial HIFα-regulated CNS angiogenesis.**

In the developing murine CNS, HIFα function in oligodendroglia (Oligo) is essential and sufficient for endothelial cell (EC) division and new vessel growth, two crucial steps of CNS angiogenesis. At the molecular level, HIFα stabilization in Oligo activates VEGF but not Wnt expression. Mechanistically, oligodendroglial HIFα modulates vascular EC properties by secreting and activating VEGF but not Wnt signaling pathway. In addition to oligodendroglia, astroglia (Astro) also regulates CNS angiogenesis and this regulation is at least in part through HIFα-activated Wnt signaling.

**Supplementary Table 1 - Primary antibodies used for immunohistochemistry in the study.**

| **Antibody targets** | **Catalog #, RRID, dilution used, and Provider** |
| --- | --- |
| EYFP/GFP | #06-896, RRID: AB_310288, 1;500, Millipore |
| Laminin | #L9393, RRID:AB_477163, 1:500, Sigma |
| PECAM1 (CD31) | #sc-1506-R, RRID:AB_831096, 1:100, Santa Cruz Biotechnology |
| ERG | #97249, RRID:AB_2721841, 1:200,Cell Signal Technology |
| PKM2 | #4053, RRID:AB_1904096,1:200, Cell Signal Technology |
| Albumin | #A90-134A, RRID:AB_67016, 1:200, Bethyl Laboratories |
| HIF1α | #NB100-105,RRID:AB_1001154,1;200, Novus |
| GFAP | #Z0334,RRID:AB_10013382, 1:400, Agilent Technologies |
| NeuN | #MAB377, RRID:AB_2298772,1:200, Millipore |
| S100β | #ab66028,RRID: AB_1142710, 1:200, Abcam |
| Sox10 | #sc-17342, RRID: AB_2195374, 1:100;Santa Cruz Biotechnology |
| BrdU | #sc-70441, RRID: AB_1119696, 1:100; Santa Cruz Biotechnology |
| Isolectin B4 (IB4) | #L2140, RRID: AB_2313663, 1:100, Sigma |
| Active β-catenin | #05-665, RRID: AB_309887, 1:200, Millipore |

**Supplementary Table 2 - Primary antibodies used for Western blot in the study.**

| **Antibody targets** | **Catalog #, RRID, dilution used, and Provider** |
| --- | --- |
| β-actin | #3700, RRID: AB_2242334, 1:5,000, Cell Signaling Technology |
| PKM2 | #4053, RRID:AB_1904096, 1:2000, Cell Signaling Technology |
| HK2 | #2867, RRID:AB_2232946, 1:2000, Cell Signaling Technology |
| MCT1 | #sc-50325, RRID:AB_2083632, 1:2000, Santa Cruz |
| β-catenin | #610153, RRID: AB_ 397554, 1:1000, BD Pharmingen |
| Active β-catenin | #05-665, RRID: AB_309887, 1:1000, Millipore |
| Axin2 | #6163, RRID: AB_10904353, 1:1000, Prosci |
| ERG | #97249, RRID: AB_2721841, 1:2500, Cell Signaling Technology |
| Naked1 | #2262, RRID:AB_561471, Cell Signaling Technology |

**Supplementary Table 3 – Primer sets of RT-qPCR used in the study.**

| ***Gene*** | **Forward primers (5’-🡪3’)** | **Reverse primers (5’-🡪3’)** |
| --- | --- | --- |
| *Vhl* | CTCAGCCCTACCCGATCTTAC | ACATTGAGGGATGGCACAAAC |
| *Wls* | AGGGCAAGGAAGAAGGAGAG | ATCCCTCCAACAATGCAGAG |
| *Pecam1* | ACGCTGGTGCTCTATGCAAG | TCAGTTGCTGCCCATTCATA |
| *Glut1* | CAGTTCGGCTATAACACTGGTG | GCCCCCGACAGAGAAGATG |
| *Pkm2* | GCCGCCTGGACATTGACTC | CCATGAGAGAAATTCAGCCGAG |
| *Ldha* | CATTGTCAAGTACAGTCCACACT | TTCCAATTACTCGGTTTTTGGGA |
| *Hk2* | TGATCGCCTGCTTATTCACGG | AACCGCCTAGAAATCTCCAGA |
| *Mct1* | TGTTAGTCGGAGCCTTCATTTC | CACTGGTCGTTGCACTGAATA |
| *Axin2* | AACCTATGCCCGTTTCCTCTA | GAGTGTAAAGACTTGGTCCACC |
| *Naked1* | CAGCTTGCTGCATACCATCTAT | GTTGAAAAGGACGCTCCTCTTA |
| *Notum* | GGACAGCTTTATGGCGCAAG | TCACCGACGTGTTCAGCAG |
| *Sp5* | GTACTTGCCATCGAGGTAG | GGCTCGGACTTTGGAATC |
| *LacZ* | CGCTGACGGAAGCAAAACA | GCCCGGATAAACGGAACTG |
| *Wnt7a* | CGACTGTGGCTGCGACAAG | CTTCATGTTCTCCTCCAGGATCTTC |
| *Wnt7b* | CTTCACCTATGCCATCACGG | TGGTTGTAGTAGCCTTGCTTCT |
| *Vegfa* | GCACATAGAGAGAATGAGCTTCC | CTCCGCTCTGAACAAGGCT |
| *Hsp90* | AAACAAGGAGATTTTCCTCCGC | CCGTCAGGCTCTCATATCGAAT |
